# Parallel G-quadruplex folds via multiple paths involving G-tract stacking and structuring from coil ensemble

**DOI:** 10.1101/2023.09.09.556957

**Authors:** Pavlína Pokorná, Vojtěch Mlýnský, Giovanni Bussi, Jiří Šponer, Petr Stadlbauer

**Affiliations:** Institute of Biophysics of the Czech Academy of Sciences, Královopolská 135, Brno 61200, Czech Republic; Scuola Internazionale Superiore di Studi Avanzati (SISSA), via Bonomea 265, Trieste 34136, Italy

## Abstract

G-quadruplexes (G4s) are non-canonical nucleic acid structures that fold through complex processes. Characterization of G4 folding landscape contributes to comprehending G4 roles in gene regulation but is challenging for experiments and computations. Here we investigate the folding of a three-quartet parallel DNA G4 with (GGGA)_3_GGG sequence using all-atom explicit-solvent enhanced-sampling molecular dynamics (MD) simulations. We suggest an early formation of guanine stacks in the G-tracts, which behave as semi-rigid blocks in the folding process. The parallel G4 folding is initiated by the formation of a collapsed compact coil-like ensemble. Structuring of the G4 from the coil then proceeds via various cross-like, hairpin, slip-stranded, and two-quartet ensembles and can bypass the G-triplex structure. While parallel G-hairpins are extremely unstable when isolated, they are more stable inside the coil structure. Folding of parallel G4 does not appear to involve any salient intermediates and, instead, it is an extremely multiple-pathway process. On the methodology side, we show that the AMBER DNA force field predicts the folded G4 to be less stable than the unfolded ensemble, uncovering substantial force-field issues. Overall, we provide unique atomistic insights into the folding landscape of parallel-stranded G4 but also reveal limitations of the state-of-the-art MD techniques.

## INTRODUCTION

G-quadruplexes (G4s) are non-canonical nucleic acid structures formed by guanine-rich sequences. Localized dominantly in telomeres and gene regulatory regions, they may contribute to genome replication and maintenance, gene expression control, and are implicated in the development of diseases such as cancer (1–3). Besides their biological roles and therapeutic targeting potential, they also find applications in nanotechnology (4, 5).

The core of G4 structure is formed by four guanine bases H-bonded together in a planar arrangement, i.e., quartet (***Figure 1***). Some DNA G4s are structurally polymorphic and can adopt several different folds with diverse orientations of their G-strands and the connecting loops (6–8). Different folds formed by the same sequence then differ in their stabilities under varying environmental conditions, such as salt conditions or protein binding (9). The *syn* (*s*) and *anti* (*a*) orientations of individual guanine glycosidic torsion angles are a basis for the *syn-anti* pattern of the whole G4. However, not all *syn-anti* patterns can form a G4; the basic rule is that if two G-strands run parallel to each other, their two guanines participating in the same quartet must both have the same glycosidic orientation. In contrast, for antiparallel G-strands two guanines in a given quartet have opposite glycosidic orientations (***Figure 1***) (10). Additionally, a single-nucleotide loop predominantly folds into a propeller topology and thus only supports a parallel G-strand orientation (11–13).

**Figure 1:**
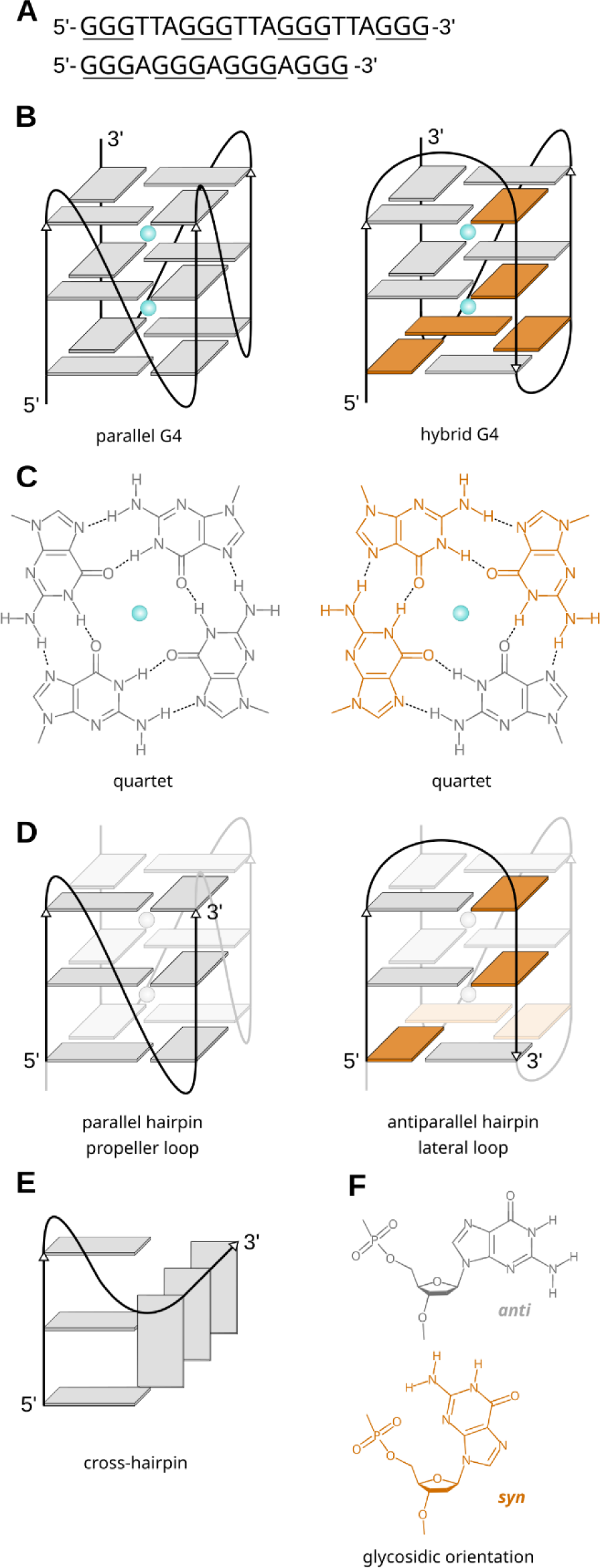
Examples of G4 topologies. (**A**) Examples of G4-forming sequences, G-tracts are underlined. (**B**) Schemes of three-quartet parallel and hybrid G4s, bases in *anti*-conformation are in grey, *syn* in orange and the blue spheres represent cations. (**C**) Structural formulas of the bottom quartets of G4s from (**B**), H-bonds are marked with dashed lines. (**D**) Hairpins derived from parallel and hybrid G4s. The propeller loop is also called the double-chain-reversal loop. (**E**) Scheme of a cross-hairpin structure. The G-tracts are perpendicular to each other. (**F**) *Anti* and *syn* guanosine orientations.

Globally, G4 folding is a complex multi-pathway process where equilibrium is often reached slowly. *In vitro* experiments revealed folding times ranging from sub-seconds up to many hours, going far beyond timescales relevant to many cellular processes (14–16). At the same time, both excessive stabilization and destabilization of G4s were reported to have pathological consequences, suggesting that the balance of folding-unfolding kinetics is important for G4 regulatory functions (17). Thus, not just the equilibrium ensembles and significant local minima G4 structures, but also pathways and kinetics of G4 folding and transitions among the folds are expected to be biologically relevant (14, 15, 18).

From the folding landscape theory point of view, the G4 folding can be best described by the kinetic partitioning mechanism, which is a competition between two or more long-living species (19). It has been suggested that the long-living species forming the dominant free energy basins on the landscape are diverse G4 folds (19, 20). The folding landscape of a G4 may contain several competing quadruplex topologies including G4 structures with formal strand slippage (register shift, reducing the number of G-quartets). For many sequences, competition between several G4 structures was experimentally detected, while other undetectable G4 species may be present transiently during the folding process. Indeed, for the most widely-studied G4, the human telomeric (GGGTTA)_n_ sequence, a multi-pathway branched folding processes involving several species with differing lifetimes was reported with NMR, circular dichroism, FRET, and mass spectroscopy experiments (18, 20–22). Obviously, the landscape must also include transitions between the different long-living species. Thus, besides diverse G4 topologies, a rich spectrum of structures and substates is expected to be transiently populated during individual folding events or transitions between different G4 topologies. Among them, G-hairpins, G-triplexes, and cross-hairpin structures (*Figure 1*) were suggested as prominent (22–33). Experimental investigations of transiently populated partially folded species are very challenging due to their short lifetimes, low populations, and structural fluxionality. Nevertheless, these structures or ensembles are important, as they can form bottlenecks for transitions between folded and unfolded states or between long-living G4 species.

MD simulations are a viable tool to study the transient ensembles and dynamic processes thanks to their detailed spatial and temporal resolution. They were used multiple times to address different aspects of G4 folding (23, 26, 27, 30, 32–47). Standard MD is limited by affordable sampling, so diverse enhanced sampling methods are often used. These act either generally by modifying the total energy of the system or on a specific selected low-dimensional projection (collective variable, CV) on the free energy surface (FES). The replica-exchange solute tempering (REST2) (48) falls into the first category, while metadynamics (49) is in the second category. We have previously used REST2 simulations to investigate all-*anti* G-sequences capable of forming parallel G-hairpins and triplexes (32). The work suggested that the perpendicular cross-hairpin orientation of the G-tracts is more stable than the ideal Hoogsteen-paired hairpin and that the parallel triplex ensemble can be one of the possible transitory folding states but is not stable enough to represent a major long-lived intermediate.

Here, we use MD simulations to investigate transitory species involved in the folding of parallel-stranded G4 with special emphasis on the G-hairpins which are assumed to play a role in the earliest stages of the folding. We use a combination of the REST2 scheme coupled with well-tempered metadynamics (49) (further referred to as ST-metaD method (50)) to accelerate the sampling. We simulate folding of a full G4 with (GGGA)_3_GGG sequence. Further, we characterize thermodynamic stabilities of various G-hairpin and cross-hairpin ensembles, including ensembles formed by the GGGTTAGGG human telomeric sequence.

While our primary goal was to investigate the folding landscape of the parallel G4 and various G-hairpins, we also pursued two methodological goals. Firstly, we wanted to assess the sampling capability of the ST-metaD method to characterize the folding of G-hairpins and potentially also full G4s. It was shown recently that the ST-metaD method substantially outperforms the REST2 method alone and is very efficient in characterization of short RNA hairpin loops (51). However, G4 and G-hairpins are much more challenging systems. Their free energy landscape is considerably more complex, and the G-hairpins are unstable species. Secondly, we wanted to assess the performance of the force field to capture the folding free energy of G4 and the intermediates.

Our simulations support the view that folding of full parallel three-quartet G4 (GGGA)_3_GGG starts with a formation of a compacted coil-like ensemble of the G-strands. The G4 structure then emerges from this ensemble by multiple pathways without a single salient intermediate structure. The folding events proceed through a series of incremental conformational changes via cross-like structures, hairpins, slip-stranded, and two-quartet G4 ensembles. The coil-like ensemble is suggested to be coordinating at least one monovalent cation. The simulations also suggest that the formation of guanine stacks in G-tracts (G-strands) is an important earliest phase of the folding process. The stacked G-tracts assemble into larger structures as semi-rigid blocks which simplifies the search of conformational space by the DNA chain, resembling the diffusion collision model of protein folding (52, 53). The isolated G-hairpins are not stable, but they can be supported by additional interactions in the compacted coil-like ensemble. Besides obtaining insights into the G4 folding process, we also extensively discuss limitations of the force field, as the fully folded G4 is predicted to be thermodynamically unstable.

## MATERIAL AND METHODS

### Starting and reference structures

We have performed all-atom explicit solvent MD simulations of DNA sequences <u>GGG</u>A<u>GGG</u>A<u>GGG</u>A<u>GGG</u>, <u>GGG</u>A<u>GGG</u>A<u>GGG</u>, <u>GGG</u>A<u>GGG</u>, and <u>GGG</u>TTA<u>GGG</u> starting from extended single-stranded chains. The starting structures were built by the Nucleic Acid Builder tool of AmberTools (54) as strands of a B-DNA helix with the complementary strand removed. In each simulation, the molecules were biased to fold into a specific topology (G4, triplex, or hairpin) defined by the reference structure (***Figure 2***). Throughout the text, the *syn-anti* patterns of the G-tracts will be denoted with “*s*” (*syn*) and “*a*” (*anti*) in the designation of the topology, e.g., *aaa*-*aaa* means all-*anti* G-tracts. The groove widths will be denoted with “w” for wide, “m” for medium, and “n” for narrow. “H” in the sequence notation (e.g., GGGH_n_GGG) means any nucleotide, but G.

**Figure 2:**
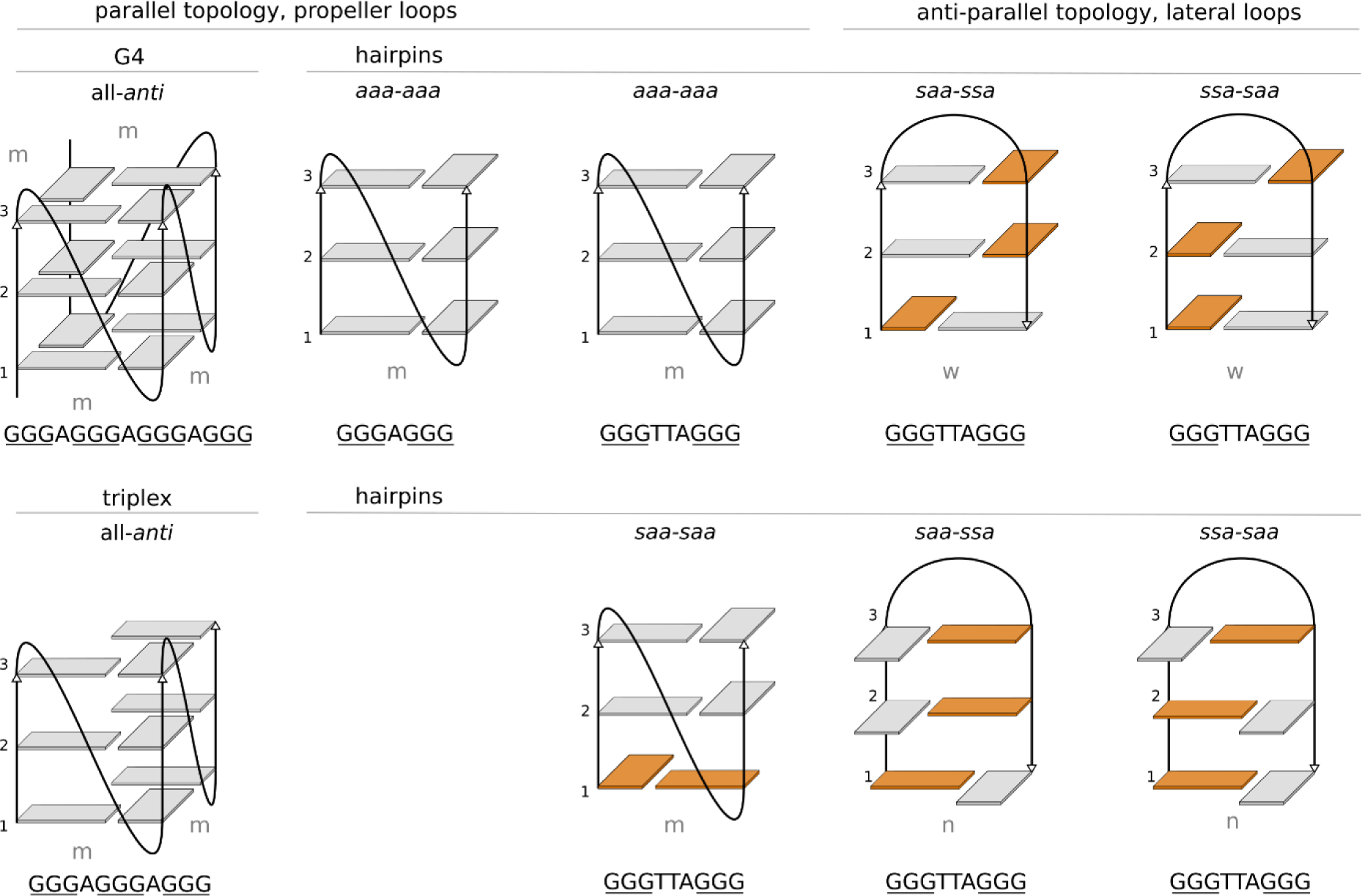
Schematic representations of the reference structures. Bases in *anti*-conformation are in grey, *syn* in orange. Letters above the hairpins denote the *syn-anti* patterns of G-tracts. Gray letters denote groove widths (wide, medium, and narrow). Sequences below the reference structures correspond to the sequences of the simulated systems.

As a reference structure for the folding of the full G4 we have used the first model of NMR structure PDB ID 2LEE (55). For the triplex and GGGAGGG hairpin, residues G3 to G13 and G7 to G13 of 2LEE were used, respectively. As reference structures for the GGGTTAGGG hairpins we used residues G2 to G10 from PDB ID 1KF1 (56) for the *aaa*-*aaa* hairpin. Nucleotides G3 to G11, G9 to G17, and G15 to G23 from PDB ID 2GKU (57) were used for the *saa-saa* hairpin, *saa-ssa* wide-grooved hairpin, and the *ssa-saa* narrow-grooved hairpin, respectively. Reference structures of the *saa-ssa* narrow-grooved hairpin and *ssa-saa* wide- grooved hairpin were derived by manual flip of the middle base pair in the corresponding hairpin taken from 2GKU, followed by proper equilibration of such a structure.

The DNA molecules were described by either the AMBER OL15 force field (58–63) or by its upgraded OL21 version (64). The molecules were solvated into octahedral boxes with SPC/E (65) water model and simulations were performed under a 0.15 M KCl salt (Joung&Cheatham parameters (66)). The minimal distance between the starting extended DNA strand and water box border was set to 25 Å for the G4 and triplex and 15 Å for the hairpin sequences, respectively. For simulations with basic loops, the charges for abasic nucleotides with hydrogen atoms attached to C1’ instead of N1/N9 were taken from ref. (67). In three simulations, we have supported guanine base-base H-bonds (N-H…N and N-H…O) with an HBfix potential (68, 69) either by stabilizing only the “native” signature H-bonds or all possible guanine base-base H-bonds generally (gHBfix; note that the gHBfix in this study included only guanine bases) (69).

### Simulation protocol

The structures were first equilibrated by minimizations with a gradual decrease of position restraints and heating. The SHAKE and hydrogen-mass-repartitioning (70) were employed to allow for a 4 ps integration time step. Then a 500 ns *nVT* simulation was run and starting structures for each replica were taken from equally distributed time intervals of that simulation. The initial velocities were randomized at the beginning of each simulation. V-rescale thermostat (71) was used with a 0.1 ps coupling-time constant to maintain the temperature of 298 K while the pressure was not regulated to keep the volume constant.

In the ST-metaD scheme, the REST2 (48) protocol runs multiple simulations in parallel and allows the individual trajectories to travel among replicas with different effective temperatures. When in higher replicas, the system can cross enthalpic barriers more easily, which facilitates conformational sampling. At the same time, bias potential along a selected collective variable (CV) is built in every replica, and its evaluation directly provides free energy profiles. In each replica, solute dihedral potentials and nonbonded interactions were scaled with a scaling factor *λ*, and solute–water interactions with 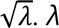 ranged from 1.00 to 0.60 for the 12 replica runs and from 1.045 to 0.600 for 16 replica runs, resulting in an effective temperature of 500 K in the highest replica. Reference replica was the one with *λ*=1.00. One simulation was performed with a broader range of 266-700 K. Replica exchanges were attempted every 10 ps and the acceptance rates ranges were between 15-50% and 20-60% for the G4 and hairpins, respectively.

The employed CVs for the ST-MetaD biasing were based on εRMSD (72), which is a form of a contact matrix describing relative orientations of bases with respect to a reference structure. εRMSD was calculated only for guanines so that the loop orientation was not included in the CV. Augmented εRMSD with a 3.2 cutoff was used for the biasing while standard εRMSD for analyses as done in ref. (51). For some simulations, an inter-tract εRMSD CV was used, where only the relative orientations of bases from different G-tracts were biased but mutual orientations of the bases within the same G-tract were not treated by this CV, to eliminate a possibility that the CV enhances stacking within the G-tracts. The CV for *N* G-tracts was defined as:

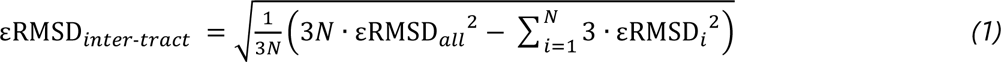

where *εRMSD_all_* is εRMSD calculated for all 3*N* guanines, while *εRMSD_i_* is εRMSD calculated only for guanines from *i*-th G-tract. The schematic representation of inter-tract εRMSD is provided in the Supplementary data, Figure S1.

Gromacs package (versions 2018 or 2021) (73) patched with PLUMED (versions 2.5.6 or 2.7.3) (74) was used to run the ST-MetaD simulations. The performed simulations are listed in Table *1*. The parameter, coordinate, and simulation settings files, reference structures, λ values, and PLUMED input files are available in the Supplementary data. The calculated bias files, reference replica ensembles, and selected demuxed trajectories were uploaded to the Zenodo database, DOI 10.5281/zenodo.8247280.

**Table 1:**
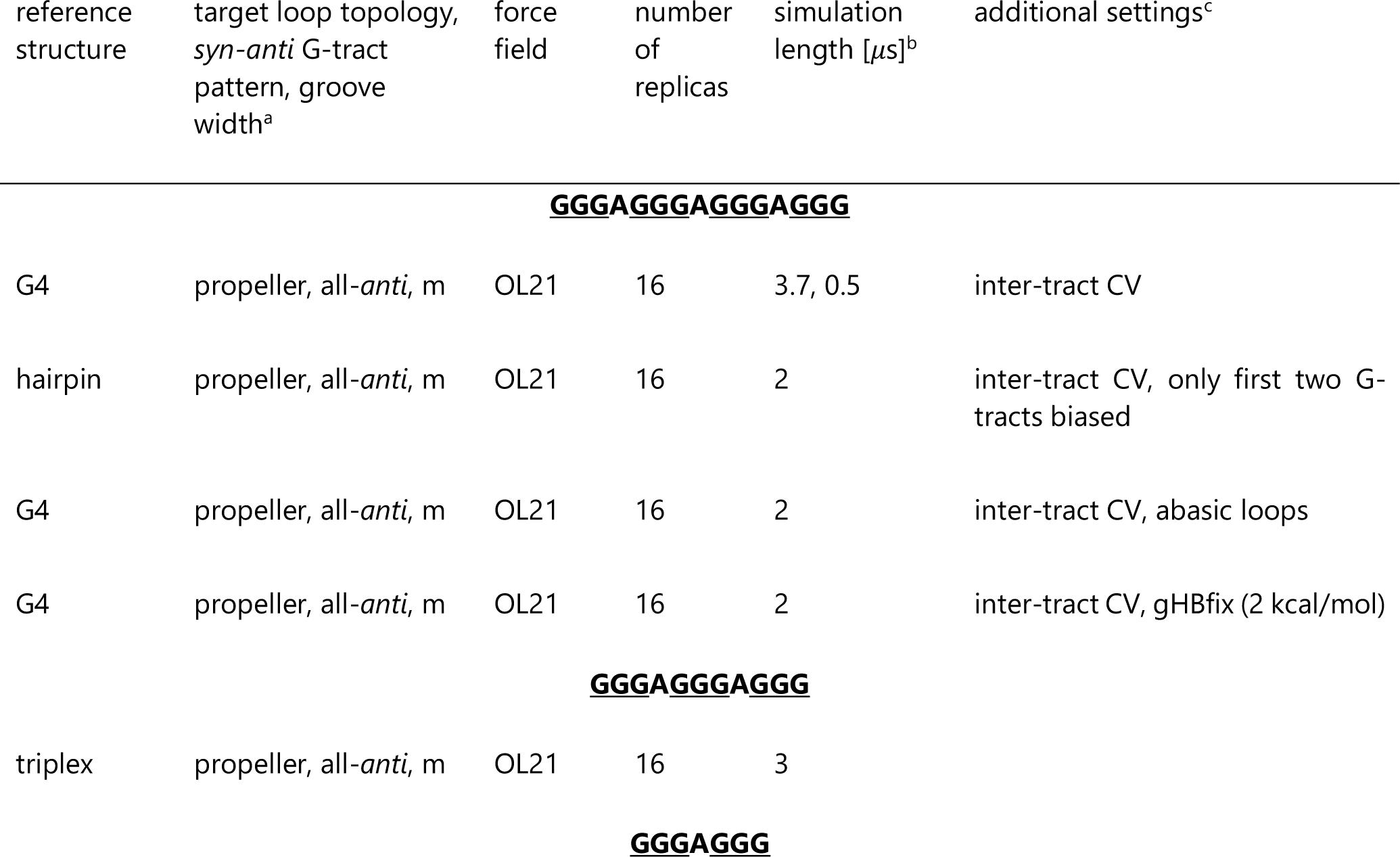
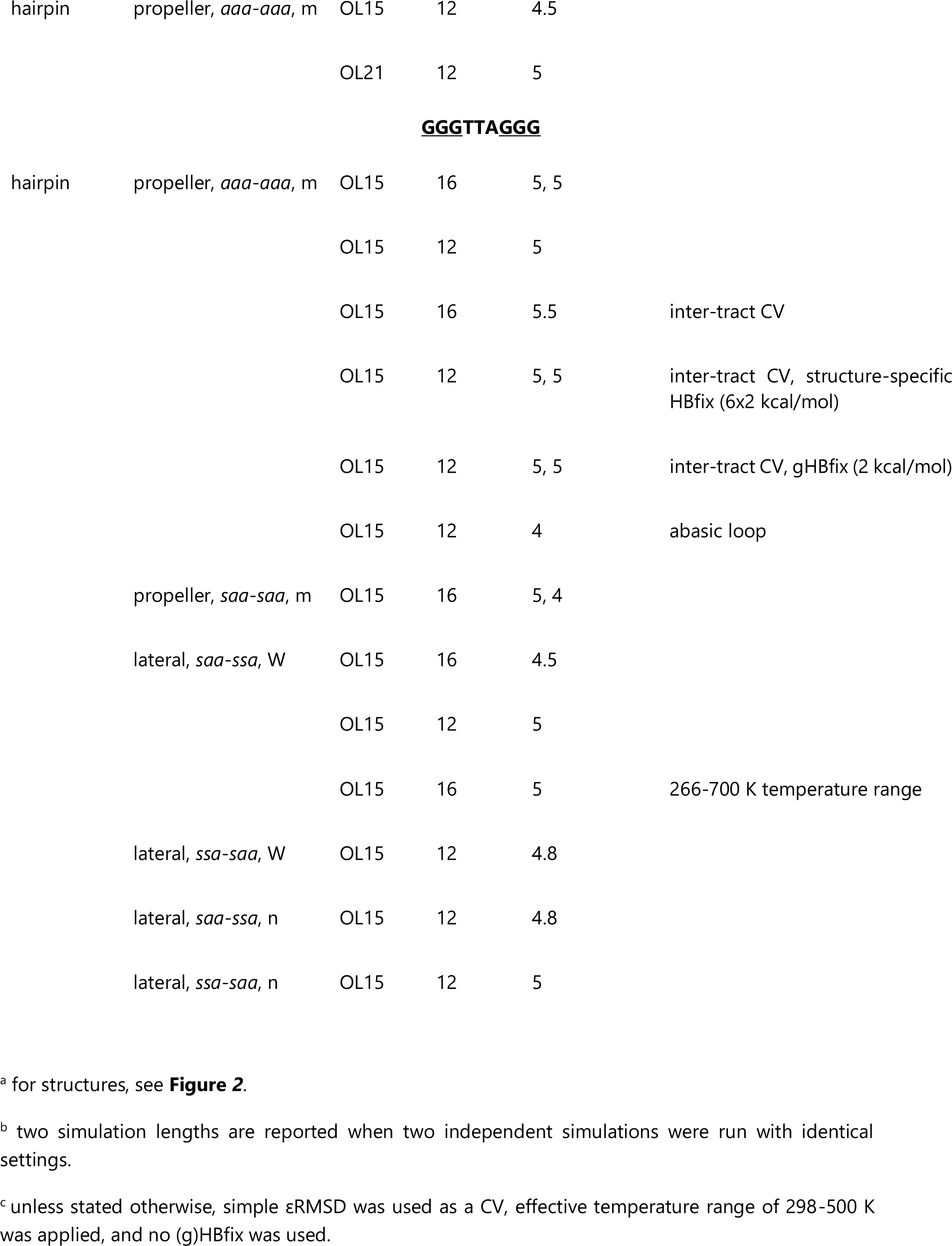
Simulations reported in this study.

### Analyses

The metadynamics bias typically converged on the timescales of a few µs as shown by ΔG_fold_ estimations in Supplementary data, Figures S2 and S3. Nevertheless, ΔG_fold_ estimations fluctuate by ∼1-2 kcal/mol over the courses of simulations, so we used time averaging of the bias potential (75) over the second half of the trajectories. All reported ΔG_fold_ values are calculated from the time-averaged bias from the reference replicas. ΔG_fold_ values from independent simulations of the same system differed by about 1 kcal/mol. We thus expect that the errors in our data are around 1 kcal/mol and in the text, we report average ΔG_fold_ values from either two or three independent simulations with the same CV (regardless the number of replicas), where applicable. Values from individual simulations are reported in Supplementary data, Table S1. Further discussion on statistical errors in similar types of calculations can be found in ref. (51).

A cutoff of 0.7 was used for εRMSD to consider two structures belonging to the same ensemble and to calculate ΔG_fold._ εRMSD was also used to identify the different structural elements (e.g., cross-hairpin or slip-stranded ensembles) in G4 and hairpin simulations.

ΔG_fold_ values for hairpin and cross-hairpin ensembles formed by the (GGGA)_3_GGG sequence were derived from the G4-biasing simulation by selecting the hairpin/cross-hairpin snapshots in the reference replica using εRMSD to the respective structures and subsequently calculating their weights based on the bias obtained from the G4 simulation. The protocol is available in the Supplementary data. ΔG_fold_ of cross-hairpin and tilted ensembles in the hairpin- biasing simulations were calculated in the same way.

While reference replica was used for ΔG_fold_ calculations, we have also monitored developments of continuous demuxed trajectories as they travel though the replica ladder to visualize the folding events.

G-tract stacking area was calculated as a sum of interaction surfaces between bases of G1 and G2, and between G2 and G3. A cutoff of 80 Å^2^ was used to consider the G-tract fully stacked. The cation coordination number (*cn*) to evaluate K^+^ binding in the G4 channel was calculated using distances between all cations and O6 atoms of guanines from two consecutive quartets (quartets *i* and *j*) as:

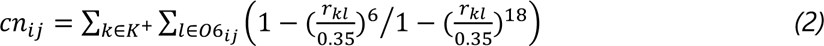

where *r_kl_*is the instantaneous distance between a pair of K^+^ and O6 atoms in nanometers.

## RESULTS

We have performed enhanced-sampling folding simulations of a parallel three-quartet G-quadruplex (G4) and of several putative G4 folding intermediates, namely various G-hairpin topologies and parallel G-triplex, to study their folding mechanisms and stabilities. The applied ST-metaD protocol with the εRMSD CV could capture hairpin and even G4 folding events. Each G4 folding event proceeded via a different route but the shared feature is the formation of the coil ensemble in the beginning. Our data further show that stacked G-tracts are important elements in hairpin and G4 folding. For isolated GGGH_n_GGG sequences, the cross-hairpin ensemble is strongly preferred over the hairpin for the all-*anti* configuration due to interactions with the loop. The force field predicts parallel hairpins with propeller loops to be notably less stable than their antiparallel counterparts with lateral loops. The predicted ΔG_fold_ for a full parallel G4 is also positive, pointing to force-field imbalances in the G4 description. In the following text, we first describe the G4 folding simulations and then the simulations of GGGAGGG and GGGTTAGG hairpins.

### Folding of the full parallel G4 proceeds via the coil ensemble

We have simulated the (GGGA)_3_GGG sequence biased to fold into a parallel G4 with single-nucleotide loops. In two simulations (3.7 and 0.5 µs long per replica) running with 16 replicas, we observed in total six G4 folding events starting from unfolded ensemble. **Figure 3** summarizes the formation of the individual structural elements of the G4 in the corresponding reactive trajectories; reactive trajectory is part of the continuous trajectory which captures the transition from the unfolded state to the fully formed state. We further observed one major subsequent refolding event involving partial G4 unfolding via strand-slippage of one G-tract and loosening of the outer quartets (**Figure 3**a bottom, trajectory 6-2). The folding events are visualized in Supplementary data, Videos 1-6.

**Figure 3:**
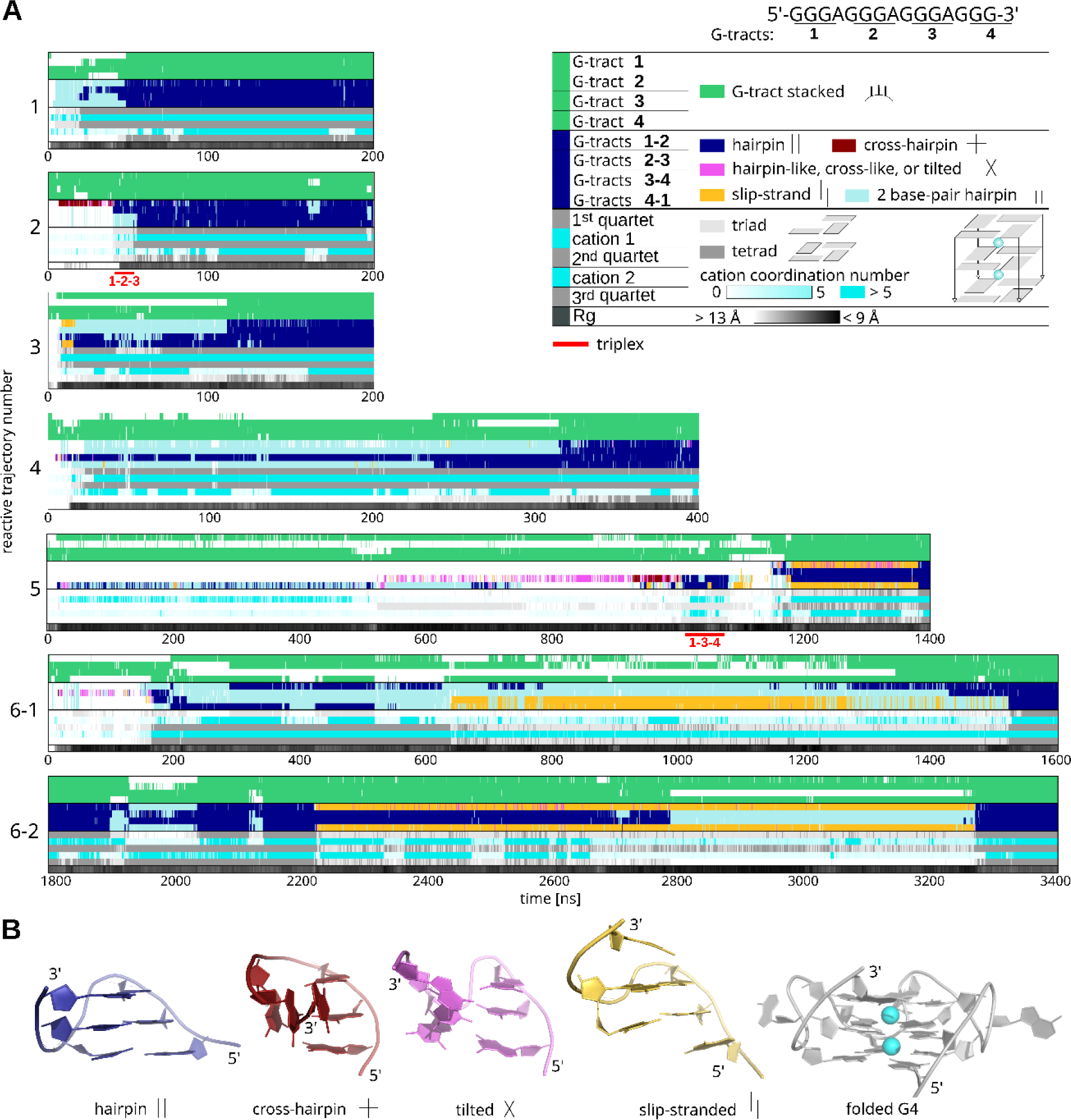
Simulated G4 folding events. (**A**) Time development of the six G4 folding events and one refolding in six reactive trajectories. Individual lines in the graphs document formations of selected key structural elements as shown in the legend. The first four lines monitor stacking in G-tracts. The next four lines monitor mutual orientations of two neighboring G-tracts; pink corresponds to a broader ensemble with εRMSD_hairpin_<1.0, εRMSD_cross_<1.0 and both G-tracts stacked, which encompasses majority of tilted structures and hairpin/cross-hairpin structures with one nucleotide in *syn*. Cross-hairpin and tilted ensembles featuring only four guanines are not plotted. The next five lines monitor the formation of quartets and cation coordination between them, and the last line monitors the compaction of the structure by its radius of gyration (Rg). Once all the colors in the graph correspond to the color panel on the left side of the legend, the G4 is fully folded. Red numbers below the triplex marks correspond to the numbers of G-tracts forming the triplex. For reactive trajectory number 6, we show events of initial folding (6-1) and subsequent refolding (6-2) in separate graphs. Timelines for full trajectories are shown in Supplementary data, Figures S4-S7. (**B**) Representative snapshots of some of the monitored structural elements formed by two G-tracts and of the fully folded G4.

An important observation is that each of the folding events proceeded via a different path. However, the first step was typically the formation of the coil ensemble with at least one ion coordinated. The coil can be viewed as a broad ensemble of compacted structures stabilized by guanine base-base H-bond interactions and was previously suggested as a possible intermediate in parallel G4 folding (19). Then we mainly observed hairpin ensembles, slip-stranded ensembles, and two-quartet G4s along the paths. Ideal cross-hairpin ensembles (as shown in ***Figure 1***e) were only formed in two folding trajectories, but the coil ensemble commonly featured cross-like states with only four, not six, guanines interacting. A variant of the cross-hairpin ensemble with fully stacked G-tracts is a tilted ensemble whose G-tract orientation is closer to the hairpin but not fully parallel. Again, the ideal tilted ensemble was seldom sampled, but the tilted-like structures with four interacting guanines were more common. The triplex structure appeared in two folding events, immediately preceding the formation of a full G4 in one of them. The other four folding events bypassed triplexes via different routes. In the last stage of the folding, when two quartets were fully formed, the third layer typically consisted of two or three guanines. The remaining G(s) would be solvent-exposed and/or stacked with the loop adenines. Once all the Gs were folded and the last quartet formed, the second cation was coordinated between the quartets as the last step. We carried out standard unbiased simulations initiated from nearly-folded snapshots (data not shown), which revealed the same chronological order – firstly, the last quartet is formed, and then the second cation is coordinated.

Since the formation of the coil ensemble could be induced by the collective variable used, we have also performed one simulation with the sequence of the full G4 but with bias only on the first two G-tracts (to fold them into a hairpin, see Methods). The folded hairpin could serve as a scaffold for the two unbiased G-tracts, but these never folded into a full G4 or at least triplex when the G-hairpin was formed. The simulation sampled a broader range of radii of gyration (Rg) values for the unbiased strand compared to the fully biased simulations, i.e., more compact as well as some more extended states were sampled (***Figure 4***a). This indicates that the coil ensemble is not induced by the CV.

**Figure 4:**
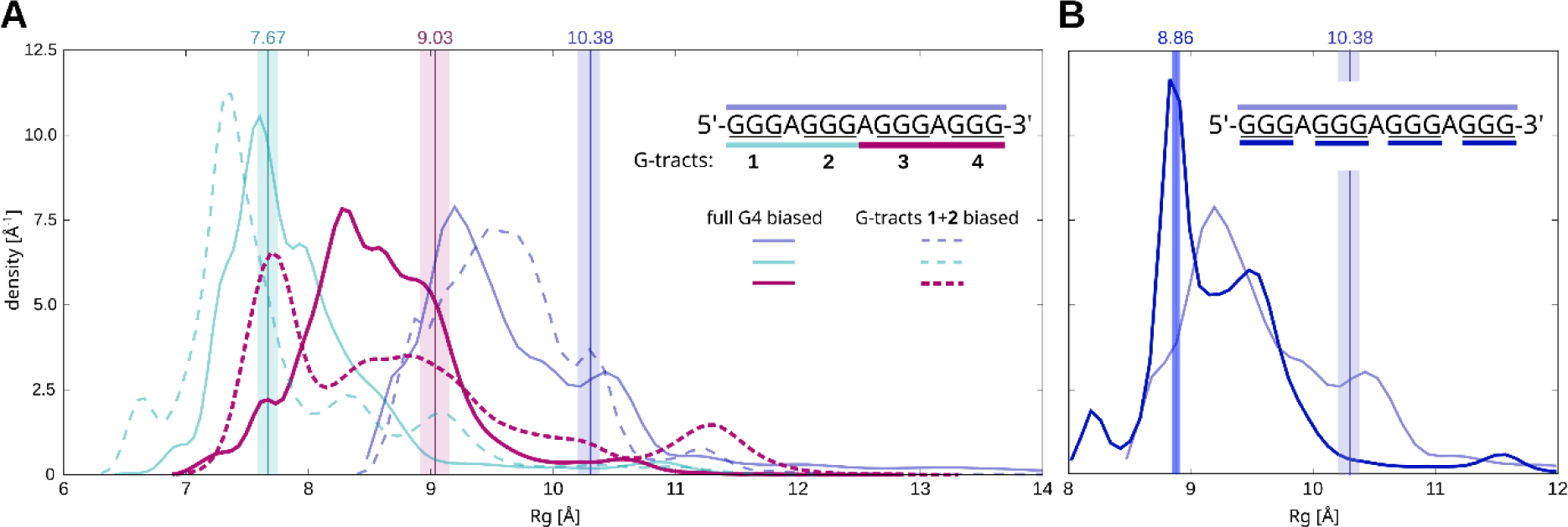
Radii of gyration in reference replicas of (GGGA)_3_GGG sequence simulations. (**A**) Values for the full G4 (violet) and its parts (GGGAGGG and AGGGAGGG, cyan and magenta) are shown with solid lines for simulations where all G-tracts were biased to fold, and with dashed lines for the simulation where only the first two G-tracts were biased. The vertical lines show median values and interquartile ranges for the folded ensemble. Reweighted densities are shown in Supplementary data, Figure S8. (**B**) Comparison of histograms for G4 calculated with and without loop adenines.

The calculation of Rg has also revealed that the exact value of overall compaction measured with radii of gyration depends strongly on the inclusion/exclusion of loop residues (***Figure 4*** b). When only guanines are considered for the calculation of Rg, the distribution peak coincides well with the native state and the majority of other states have higher Rg (***Figure 4*** b). Nonetheless, structures within the limited range around the native Rg value are not only the native G4 and closely related structures, but they also contain structures far away from the G4, as can be seen in the εRMSD distribution graph (Supplementary data, Figure S9). When Rg of the whole molecule is calculated, most of the observed structures have Rg values smaller than the native G4 (***Figure 4*** a). These structures correspond to the compacted coil ensemble. The results demonstrate that using the Rg parameter to assess the folding is an ambiguous metric.

### Force field predicts positive value of the G4 folding free energy

The ST-metaD simulation predicts a high positive ΔG_fold_ of +17 kcal/mol for the fully folded G4 (Table 2). It is evident that the force field does not properly capture the balance between folded and unfolded ensembles. To the best of our knowledge, it is the largest free energy discrepancy in simulations of nucleic acids reported in the literature so far. Despite this, we still assume that the simulations sample relevant folding trajectories and the positive free energy of the native G4 structure does not invalidate the folding events reported above. The ST-metaD method is constructed to sample high-energy states by flattening the free energy surface and thus finds the native structure despite the high positive ΔG_fold_ value and the evident large force-field imbalance over the overall FES. The force field issue will be discussed later in the paper in detail. The parallel G-hairpin has a positive ΔG_fold_ of +18 kcal/mol while the triplex is the least stable of the studied species with ΔG_fold_ of +20 kcal/mol (Table 2).

**Table 2:** Folding free energies for all-*anti* parallel G4, hairpin, and cross-hairpin ensembles.

| simulation | simulated sequence | $\Delta G_{\text{fold}}$ [kcal/mol] | | | |
| --- | --- | --- | --- | --- | --- |
|  |  | G4 | triplex | hairpin<sup>a</sup> | cross-hairpin<sup>a</sup> |
| G4 | (GGGA)<sub>3</sub>GGG | 16.9 | - | 14.4/8.3/11.1 | 12.1/5.5/8.6 |
| G4, only first two G-tracts biased | (GGGA)<sub>3</sub>GGG | - | - | 9.1/-/- | 6.0/-/- |
| triplex | (GGGA)<sub>2</sub>GGG |  | 20.1 | 8.2/6.1 | 5.7/3.0 |
| hairpin | GGGAGGG |  |  | 17.8 | 6.8 |

<sup>a</sup> calculated as $\Delta G_{\text{fold}}$ considering only hairpins (cross-hairpins) 1-2 / 2-3 / 3-4, where numbers denote the order of G-tracts in the strand (1-2-3-4), see Methods for details. "-" means that the structure was not sufficiently sampled.

Demuxed (continuous) trajectories indicated that adenine stacking stabilizes the unfolded states and slows down the folding into the complete G4. A 2-µs simulation of the full G4 with abasic loops revealed a substantially lower ΔG_fold_ of about +9 kcal/mol. This simulation also sampled more (seven) folding events. They showed principally the same mechanism as the (GGGA)3GGG simulations, proceeding through the coil ensemble via diverse pathways (Supplementary data, Figure S10).

We attempted to stabilize the G4 by generally strengthening all pairs of possible guanine base-base H-bonds (i.e., by gHBfix, see Methods). However, the extent of stabilization of the G4 fold was equivalent to the stabilization of the unfolded ensemble, so that the resulting ΔG_fold_ was within the error the same as without any force-field modification. Details are provided in the Supplementary data.

### Parallel G-hairpins are stabilized inside the coil

Cross-hairpins were previously suggested to be more stable than parallel hairpins based on their larger lifetimes in simulations (32). We have calculated the folding free energies of these structural elements from the simulations of the full G4 sequence, triplex, and the isolated GGGAGGG sequence by selecting the hairpin and cross-hairpin structure snapshots using the εRMSD to these structures (see Methods). Stabilities of the individual parallel hairpins calculated from G4 and triplex simulations are notably higher (ΔG_fold_ is less positive) than when only the isolated GGGAGGG sequence was simulated (Table 2). The stability of the cross-hairpin ensemble is comparable in both cases.

### Isolated parallel all-*anti* hairpins with propeller loops are extremely unstable while antiparallel hairpins with lateral loops are more stable

We complemented the folding simulations of the full G4 and of the isolated GGGAGGG hairpin by simulations of the isolated GGGTTAGGG sequence. In comparison to GGGAGGG, the GGGTTAGGG sequence can form a broader spectrum of hairpin topologies including antiparallel hairpins with lateral loops. While only six folding events were sampled within a cumulative time of 67.2 µs in simulations of the full (GGGA)_3_GGG G4, folding events are readily sampled in hairpin simulations (illustrated for the bigger system with TTA loop in **Figure 5**). The folding free energies for the different topologies we have targeted are provided in Table 3.

**Figure 5:**
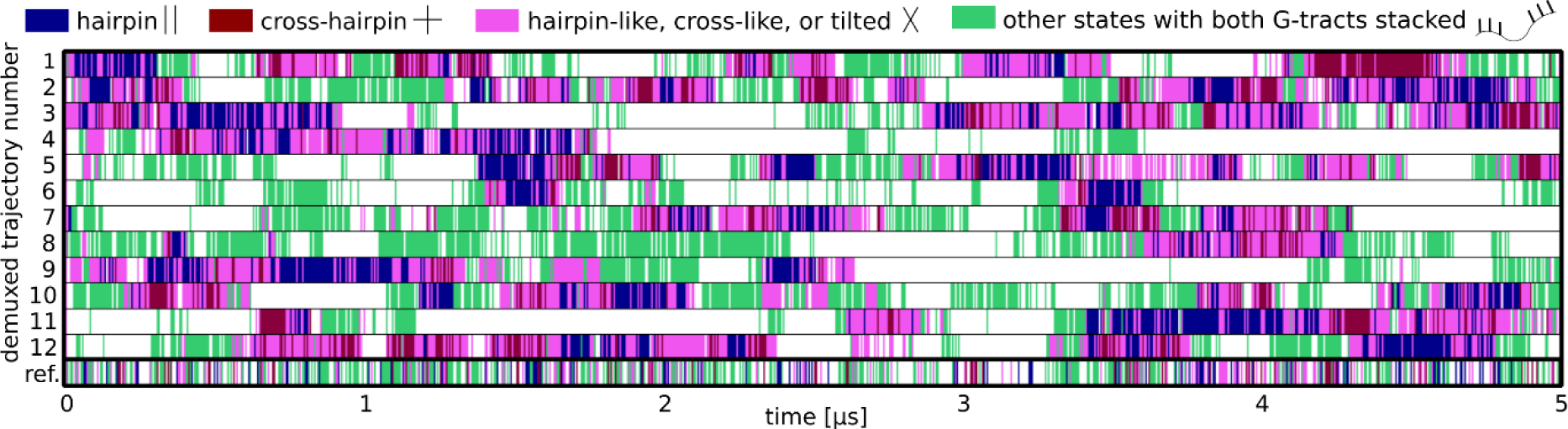
Development in twelve individual demuxed (continuous) trajectories in one GGGTTAGGG *aaa-aaa* folding simulation. Reference replica is plotted at the bottom. For details of the color-coded ensembles, see the label of **Figure 3**; green color in this figure corresponds to all other states with each G-tract having stacked guanines.

**Table 3:**
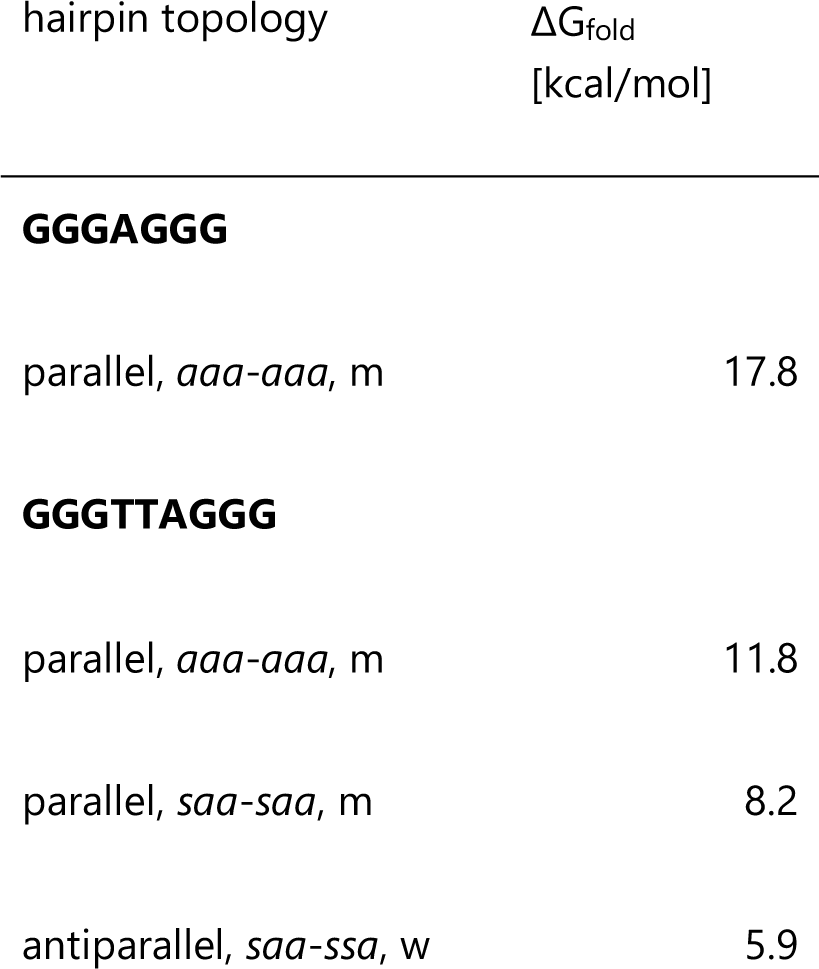
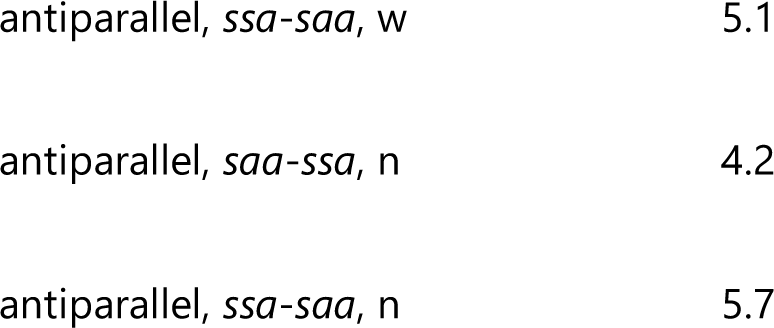
Folding free energies for the studied isolated hairpins.

| hairpin topology | $\Delta G_{\text{fold}}$<br>[kcal/mol] |
| --- | --- |
| **GGGAGGG** |  |
| parallel, *aaa-aaa*, m | 17.8 |
| **GGGTTAGGG** |  |
| parallel, *aaa-aaa*, m | 11.8 |
| parallel, *saa-saa*, m | 8.2 |
| antiparallel, *saa-ssa*, w | 5.9 |
| antiparallel, *ssa-saa*, w | 5.1 |
| antiparallel, *saa-ssa*, n | 4.2 |
| antiparallel, *ssa-saa*, n | 5.7 |

While the isolated ideally paired parallel all-*anti* GGGAGGG hairpin is extremely energetically unfavorable (+18 kcal/mol), the all-*anti* parallel hairpin formed by the GGGTTAGGG sequence has lower, yet still high, ΔG_fold_ of about +12 kcal/mol (Table 3 and ***Figure 6***). The GGGTTAGGG parallel hairpin with one *syn*-*syn* pair is more stable compared to all-*anti* (Table 3 and ***Figure 6***). This can be in part due to the G7 intra-nucleotide N2-H2…OP H-bond as suggested earlier (37). Antiparallel hairpins with lateral loops formed by the GGGTTAGGG sequence are notably more stable than parallel hairpins with propeller loops in our simulations (Table 3 and ***Figure 6***). Data for two distinct target parallel and four antiparallel hairpins clearly confirm the trend.

**Figure 6:**
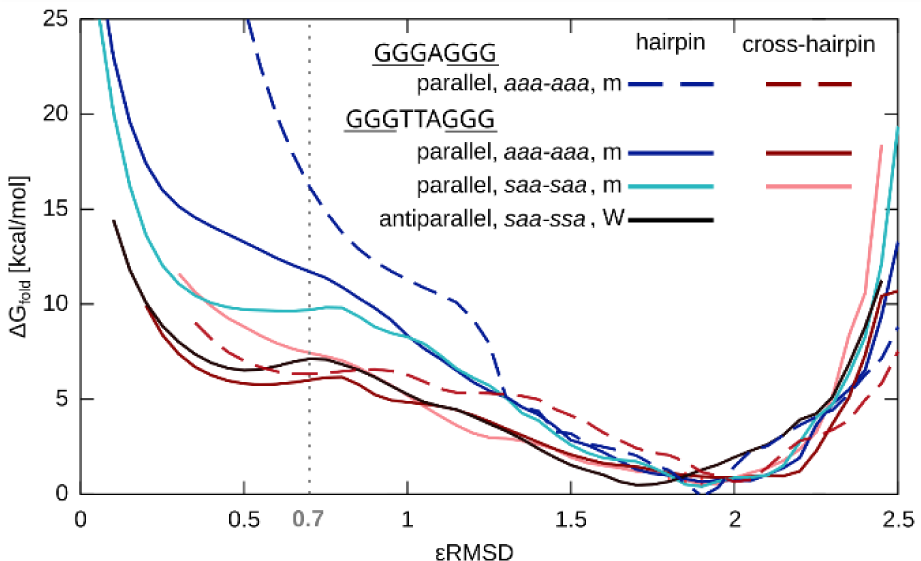
Δ_Gfold_ along the εRMSD CV for GGGAGGG and GGGTTAGGG sequences folding to different reference structures. ΔG_fold_ for cross-hairpin ensembles were calculated from the respective hairpin-biased simulations. Time-averaged bias potentials from the reference replica only were used to calculate the ΔG_fold_. The bin width used to generate the plot was 0.01.

For the case of GGGTTAGGG *aaa-aaa* parallel hairpin, we have also considered only an ensemble with all guanines in *anti* to estimate ΔG_fold_ (by removing all other snapshots from the reference replica). This allows us to effectively eliminate the competition among other possible GGGTTAGGG hairpin folds and their on-pathway states. Hairpin stability then increased by ∼3 kcal/mol, which is still above the values for antiparallel hairpins (antiparallel hairpins are stabilized by ∼1-3 kcal/mol when considering only their target *syn-anti* pattern in the ensemble). We also tried to stabilize the *aaa*-*aaa* hairpin by supporting the formation of each of the six structure-specific guanine-guanine H-bonds with 2 kcal/mol HBfix potential. Such a dramatic intervention, i.e., stabilization of the hairpin state by 12.0 kcal/mol, resulted in ΔG_fold_ of about 2.7 kcal/mol while also stabilizing some structures in the unfolded ensemble. However, stabilizing the hairpin fold did not result in an increased sampling of the folding events (Supplementary data, Figure S11). gHBfix had a smaller but opposite effect, stabilizing the unfolded ensemble (ΔG_fold_ for the hairpin was +14 kcal/mol).

### Cross-hairpin ensembles are stabilized by interactions with the loop

Cross-hairpin ensembles are more stable than hairpins for the isolated GGGH_n_GGG sequences (***Figure 6***). ΔG_fold_ values for the all-*anti* cross-hairpin ensembles of the GGGAGGG and GGGTTAGGG sequences are +6 and +5 kcal/mol, respectively. ΔG_fold_ for the tilted ensembles is about +9 kcal/mol in both cases. The cross-hairpin ensemble of the GGGTTAGGG sequence is stabilized by diverse additional stacking and H-bonding interactions with the loop residues (an example is shown in **Figure 7**a). Antiparallel hairpins are stabilized by lateral loops in a similar fashion. The GGGAGGG sequence forms lesser interactions between the G-tracts and the loop. Loop residue A4 can stack on either G3 or G5 but no specific H-bonds between the loop and the guanines are formed in the GGGAGGG cross-hairpin ensemble.

**Figure 7:**
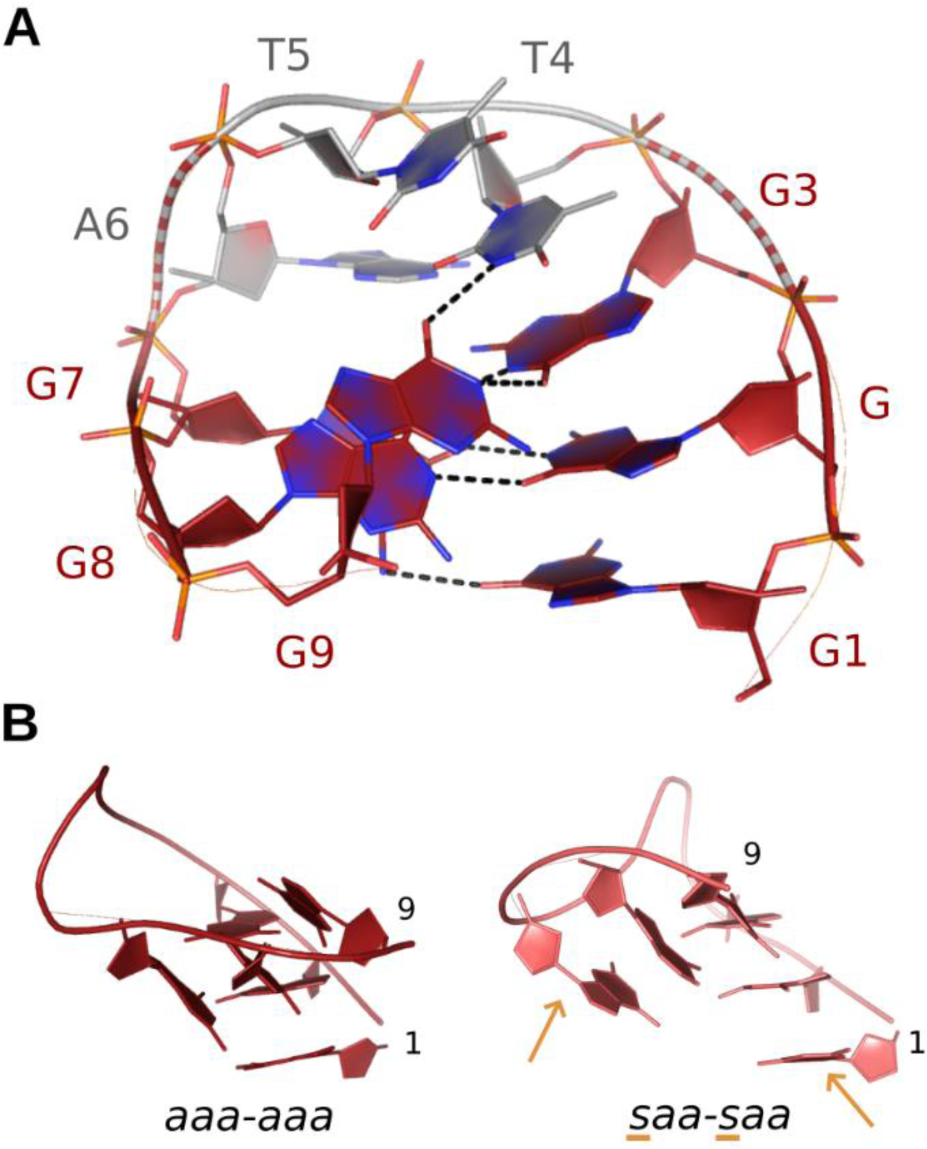
Cross-hairpin structures. (**A**) Example of loop interactions stabilizing the GGGTTAGGG cross-hairpin structure. H-bonds are marked with dashes. T4 stacks with G3 and H-bonds to G9. A6 can also stack with G7 to stabilize the cross-hairpin. (**B**) Comparison of *aaa-aaa* and *saa-saa* cross-hairpins. H-bonding with T4 was also observed in *saa-saa* cross ensemble (not shown). Bases in *syn* are indicated by orange arrows.

Solvent-exposed bases of propeller loop should introduce entropic penalty for parallel hairpins compared to loops stabilizing the folded antiparallel hairpins or cross-hairpins. Indeed, simulations with an abasic loop predicted ΔG_fold_ of about +10 and +7 kcal/mol for the GGGTTAGGG *aaa-aaa* hairpin and the cross-hairpin ensembles, respectively.

In the case GGGTTAGGG sequence and the *saa-saa* configuration, the formation of the full cross-hairpin or tilted structures with all six guanines H-bonded (as shown in **Figure 3**b for all-*anti* configuration) was not observed. The *syn*-configuration of the bases does not allow for creating compatible H-bond donor-acceptor patterns along the whole G-tract as in the case of all-*anti* configuration. When cross-hairpin or tilted structures were sampled, they had the *anti-*bases H-bonded, but one or no *syn*-base was involved in the inter-tract binding (**Figure 7**b). Thus, only five or four bases were participating in the cross, which is similar to the cross-like ensembles involving four bases seen in the G4 folding simulations. The ΔG_fold_ for the *saa*-*saa* cross-hairpin ensemble is about 1.5 kcal/mol lower compared to the hairpin, so the difference is not as pronounced as for the *aaa*-*aaa* configurations (***Figure 6***). Finally, cross-hairpin ensembles featuring a G-tract with two *syn* nucleotides were not observed.

### Hairpin formation proceeds mainly by stacked G-tracts orienting themselves with respect to each other in the space

In the demuxed folding trajectories of all systems, we commonly observe formation of G-tract stacks, i.e., the stacking of the three consecutive guanines. These stacked G-tracts then searched the space to form the target hairpin eventually (**Figure 5**). The chosen CV could drive the formation of the G-stacking in G-tracts in our hairpin-folding simulations as we targeted the relative orientations of guanine bases (described with εRMSD, see Methods). Thus, we performed one additional hairpin simulation also with the modified inter-tract εRMSD CV (see Methods for details) that does not affect mutual orientations of guanines within G-tracts. G-tract stacking was only slightly less sampled, and the stacked G-tracts were still clearly formed (Supplementary data, Figure S12). Analysis of our previously published REST2 trajectories (32) gave about 75% population of each individual G-tract fully stacked for the GGGAGGG simulation, and the number was 40% for GGGTTAGGG (Supplementary data, Table S2). Formation of stacking within the G-tracts thus seems relevant for the real folding mechanisms. We still caution that base stacking has been suggested to be overestimated by the AMBER force field (76–78), so that the magnitude of GGG stacking in our simulations may be somewhat overestimated by the force field.

### Sampling problem on the rich FES

In general, multiple folding events were observed in hairpin simulations though some demuxed trajectories did not sample hairpin folding events at all (Supplementary data, Table S1 and Figure S11). Convergence of the sampling in individual trajectories was even poorer in G4 simulations. This is not surprising given the rich structural dynamics of the studied systems and their high free-energy instability. The individual demuxed trajectories sufficiently sampled all replicas in the ladder (Supplementary data, Figure S13). Other than the target (biased) hairpin topology was rarely sampled in simulations, despite no restraints on glycosidic or backbone torsions being applied. For example, the parallel GGGTTAGGG *saa*-*saa* hairpin is sampled in 0.06% snapshots in the simulation targeting the *aaa*-*aaa* hairpin while not at all in the simulations targeting the lateral loops. The very unstable *aaa*-*aaa* hairpin is not sampled in the *saa*-*saa* simulation, but the *aaa*-*aaa* cross-hairpin is. For the more stable antiparallel hairpins, both wide and narrow groove topologies with the same *syn*-*anti* patterns are easily sampled in simulations, despite only one of them being targeted in a particular simulation. Sampling of both types of hairpins with the same *syn*-*anti* pattern in one simulation can be supported by the CV biasing the G-tracts to the same respective glycosidic patterns. The wide and narrow groove topologies with the same glycosidic patterns differ in εRMSD by ∼1.4.

The sampling issue is also documented by very limited overlap of the unfolded ensembles of simulations targeting different hairpin topologies; see Supplementary data for details. On the other hand, changes in the simulation protocol (number of replicas, different scaling factors) do not affect the results (Supplementary data, Table S1 and Figure S11). It indicates that the methodology is robust enough, or at the limits of what can be achieved with contemporary methods and computers.

## DISCUSSION

We have carried out a series of ST-metaD enhanced-sampling folding simulations targeting a three-quartet parallel-stranded DNA quadruplex (GGGA)_3_GGG and several isolated G-hairpin topologies. The goal was to better understand G4 folding, the roles of selected intermediates and, mainly, to visualize transient structures. We wanted to predict types of structures which may occur along the folding pathways; for this goal the ST-metaD method should be sufficiently robust. It is obvious that to capture complete G4 folding using unbiased MD simulations is presently entirely beyond the capabilities of contemporary computers, as well as force fields.

The folding simulation of (GGGA)_3_GGG G4 predicts ΔG_fold_ of +17 kcal/mol. This is evidently in sharp disagreement with experiments and shows the magnitude of the overall imbalance in the force field (see below). Nevertheless, the simulations in our opinion still provide useful insights into the folding mechanism of the all-*anti* parallel-stranded G4. The metadynamics method flattens the free energy landscape along the chosen CV and thus allows to sample folding events even in case of positive free energy of the target structure. The simulations show that the folding of parallel-stranded G4 is a multi-pathway process involving the structuring of quartets inside the compact coil ensemble.

When considering ideal Hoogsteen-paired structures, the isolated parallel all-*anti* G-hairpins are predicted to be strikingly unstable with ΔG_fold_ of +18 and +12 kcal/mol for the GGGAGGG and GGGTTAGGG sequences, respectively (Table 3). As discussed below, these high instabilities may partly reflect inaccuracies of the simulation force field. Formation of cross-hairpin ensembles that are on pathway ensembles to the parallel G-hairpin is a more likely event for isolated sequences. Antiparallel GGGTTAGGG G-hairpins are predicted to be more stable with ΔG_fold_ ranging from +4 to +6 kcal/mol. The simulations reveal very rich unfolded ensembles for the studied sequences, with a plethora of competing states, and do not indicate an existence of a single stable structure for the hairpin sequences.

### Suggested properties of the parallel G4 folding landscape

Despite the evident force field and sampling limitations, we suggest that the observed folding events provide useful insights into the nature of folding pathways and transitory ensembles sampled during the folding. Our simulations reveal that the conformational landscape of G4 sequences is extremely rich.

The suggested folding process of a parallel G4 (assuming entirely unstructured starting ensemble) based on the simulations is summarized in ***Figure 8***. First, extended G-tract chains form stacked G-tracts with occasional fraying of one of the three stacked guanines. These tracts then explore the conformational space and position themselves with respect to each other in many possible ways. They tend to assemble into larger structures as semi-rigid blocs. This simplifies the search of conformational space by the DNA chain, resembling the diffusion collision model of protein folding with prefolded α-helices. The parallel G4 folding proceeds from extended chains via a formation of the coil ensemble. The coil can be viewed as a broad ensemble of compacted structures stabilized by guanine base-base H-bond interactions. The coil ensemble coordinating at least one cation has been previously suggested as a folding intermediate for parallel G4s in computational and experimental studies (14, 22, 23, 32, 79). The present simulations further support this hypothesis. The compacted coil ensemble can perhaps be likened to the molten globule state in protein folding. Despite all collapsing initially into the coil ensemble, each of the six simulated G4 folding events proceeded via a different path (**Figure 3**a). Thus, the observed folding events do not represent a simple pathway through some salient intermediates. Instead, literally, the final quadruplex structure grows up from the compacted unfolded ensemble via numerous incremental conformational changes. Native interactions are formed from inside the coil and the ensemble diffuses via different possible orientations of the G-tracts, cross-like, hairpin, and slip-stranded structures until a G4 is formed. We further support the view that the triplex is not a mandatory intermediate on parallel G4 folding landscape and can be easily bypassed. Occasional events of coil untying and partial chain extension will allow for major structural rearrangements. Indeed, such chain extension events were sampled in the partially biased G4 simulation as illustrated by the range of compactness in ***Figure 4***a. The (GGGA)_3_GGG sequence can, in theory, also form an antiparallel Hoogsteen-paired hairpin featuring only two G-pairs and a two-nucleotide lateral loop. Such species did not occur in our simulations likely due to the limited sampling, but we expect them to be present on the parallel G4 folding landscape, which is also possibly consistent with some experimental observations (22, 79).

**Figure 8:**
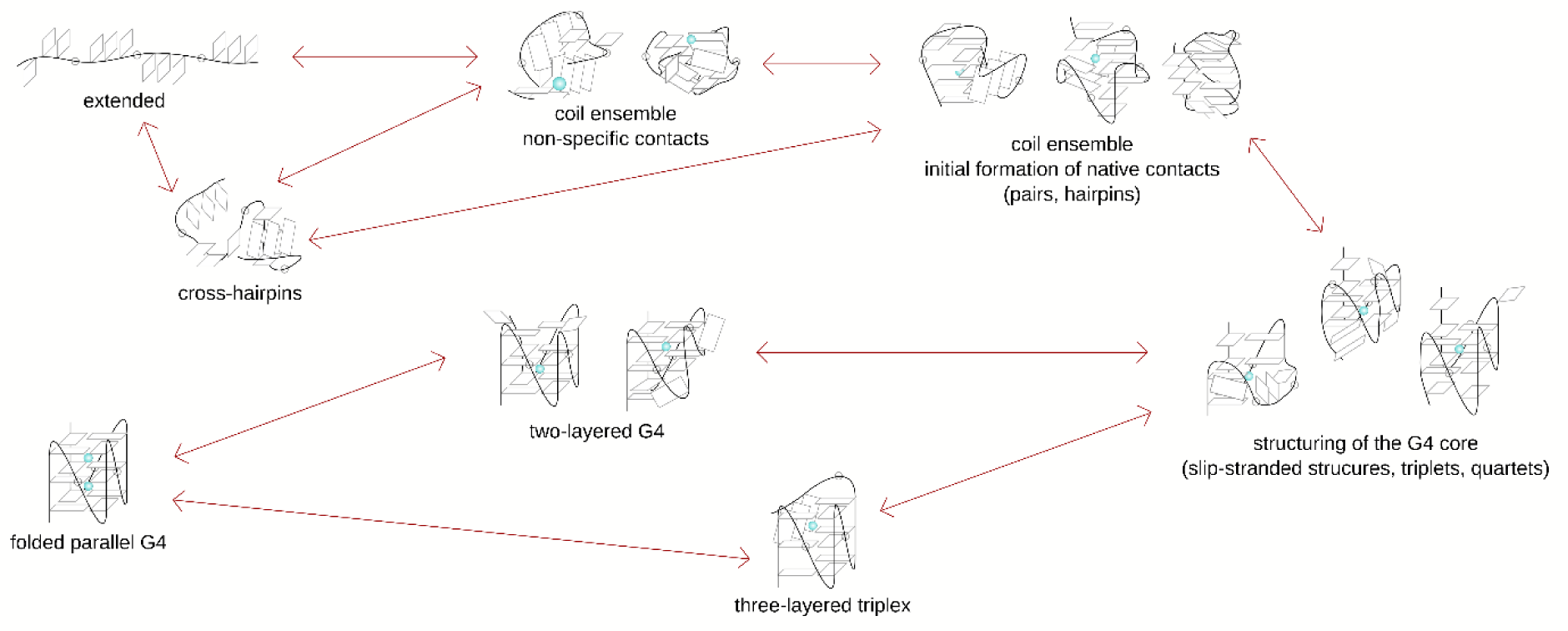
Summary of the single-nucleotide-loop parallel G4 folding process based on the presented simulations. The folding is initiated by the formation of the compact coil structure inside which the native interactions form gradually. Loop nucleotides are represented by circles on the backbone. They can also participate in stacking interactions with the guanines. The structures shown are just representatives of the broader ensembles they are parts of. Lateral species are not included in the scheme though we expect them to be present in the early-stage coil ensemble as well. We would like to point out that although we suggest that the ST-metaD simulations can quite reliably disclose types of transitory structures that participate in the folding, they do not provide any kinetic information. The depicted intermediates are not necessarily stable species and may belong to the transition ensembles.

A similar folding mechanism could also apply for parallel-stranded RNA G4. The compacted coil-like ensemble we have predicted here for the full DNA G4 may possibly be related to the hydrophobic collapse suggested by experiments for the initial phase of folding of RNA G4s (80), and so we find similarities with the multi-pathway gradual rise of RNA G4 from cross-like intermediates suggested by MD simulations (31).

Loop conformational degrees of freedom add another layer of complexity to the folding process. Simulations with abasic loops showed notably lower ΔG_fold_ for the G4 and a decrease of the ΔG_fold_ difference between the *aaa*-*aaa* hairpin and the cross-hairpin, confirming the effect of the loop residues on their stabilities. Consistently, stabilization of G4 fold upon removing loop bases was reported for single-nucleotide G4 with melting experiments (11). This could result from the elimination of adenine-stabilized unfolded structures as well as the reduction of the entropic penalty introduced by the otherwise solvent-exposed adenine base in the folded G4 (11, 81).

A single isolated hairpin sequence has the properties of a disordered chain and transits among a plethora of competing states. Spontaneous formation of a parallel hairpin without external stabilizing interactions is notably less likely than the formation of antiparallel hairpins and cross-hairpins, which are stabilized by the loop residues (**Figure 6**, Table 3). Cross-hairpin ensembles can be formed by *aaa*-*aaa* and, to some extent, *saa*-*saa syn-anti* patterns. The cross-hairpin ensemble can lead to a parallel hairpin upon rotating the relative orientations of the two G-tracts. The relative difference in stabilities of parallel hairpin and cross-hairpin decreases when the sequence is embedded in a larger context as shown by our simulations of the parallel G4 and triplex (Table 2). Residues preceding and following the GGGAGGG motif and interactions inside the G4 coil ensemble can stabilize the parallel hairpins. This is consistent with our previous simulations of G-triplex, which was stabilized by flanking nucleotides or stacking with hemin ligand (32). Thus, cross-hairpin structures may be more relevant than hairpins for the initial phases of the folding when the DNA strand is relatively extended, and individual G-tracts approach each other (initial phases of nucleation of the coil). The hairpins would be later structured inside the coil. While the cross-hairpin ensemble is suggested to dominate over the hairpin for the all-*anti* configuration in an extended (uncompacted) chain, the difference is not as pronounced for the *saa-saa* configuration. Less likely transition to the cross-hairpin might be one of the reasons for higher stability of *saa-saa* hairpin compared to *aaa-aaa*.

Given the apparently different properties of putative on-pathway folding intermediates for different G4 topologies (e.g., parallel and antiparallel hairpins studied here), the suggested folding mechanism cannot be simply generalized outside the context of the studied G4. In fact, even similar G4s can follow distinct folding mechanisms (79). Our data show that parallel hairpins need to be stabilized inside the coil, but antiparallel ones can probably form easier and in earlier stages of folding events. While we suggest that initiating the folding of a parallel G4 with one parallel hairpin formed and the rest of the chain extended is unlikely, we speculate that such a scenario can be possible for G4s containing antiparallel hairpins. Yang et al. previously reported such a folding pathway for a two-quartet antiparallel G4 of thrombin-binding aptamer (15-TBA) with lateral loops (30). Their simulation data suggested that the folding proceeds first through a hairpin-like base-collapsed ensemble where one native lateral loop is already formed but the rest of the bases are paired non-specifically and the chain then eventually structures into a triplex. The last step was binding of the last GG tract, either directly or via a strand slip. Bian et al. combined all-atom simulations with a structure-based coarse-grain model to study the folding of human telomeric G4 sequence into two different hybrid topologies (47). The first folding step predicted is the formation of an antiparallel hairpin connecting the 5’ and 3’-end G-tracts. The two topologies studied by Bian et al. folded with paths with different complexities and only one of them transited via triplex ensemble.

### Force field struggles to accurately capture the global G4 folding landscape

Though the present simulations provide unique insights into the G4 folding landscape, they are obviously limited by the force-field accuracy, selected CV, and achievable sampling. The most visible illustration of evident force field issues is the ΔG_fold_ +17 kcal/mol of the (GGGA)_3_GGG G4. In comparison, ΔG_fold_ for the (GGGA)_3_GGG G4 estimated from melting curves at 1 mM KCl solution is about -1.4 kcal/mol (11). The exact value depends on the experimental setup, but the studied G4 should have a negative ΔG_fold_. Since our simulated unfolded ensembles certainly still do not cover the whole real space of unfolded ensembles, the predicted positive ΔG_fold_ could even be underestimated. It might seem to be a surprise since standard simulations of folded cation-stabilized quadruplexes are very stable, so quadruplexes have usually been assumed to be well-described by the force field (19). Apparently, this is not true for their global free energy landscape.

Positive folding value (albeit much milder) with AMBER force field has been reported earlier by Yang et al. for the 15-TBA (30). Yang et al. suggested that H-bonding in quartets is understabilized by the AMBER force field non-bonded terms. They have proposed a correction based on a modification of van der Waals parameters using the non-bonded fix (NBfix) approach, which increased the stability of 15-TBA by ∼6.5 kcal/mol (from +3.8 kcal/mol to - 2.8 kcal/mol), i.e, by ∼3 kcal/mol per quartet. Simple general (not structure-specific) force field correction increasing guanine base-base H-bond strength (gHBfix) did not improve (GGGA)_3_GGG G4 stability in our study. The balance of H-bond strengths in quartets and in the unfolded ensemble could still be one of the sources of the positive folding energy, but the understabilization of (GGGA)_3_GGG is evidently much larger compared to the 15-TBA.

Another known force-field issue is the overestimation of ion-ion repulsion in the G4 channel due to missing polarization/charge transfer effects with fixed point charges force fields (82). Our folding trajectories suggest that binding of the second ion is the very last step in the folding after all the quartets are already fully assembled. It is possible though, that binding of the second ion proceeds faster and in earlier stages of the folding. It is likely that over- estimation of ion-ion repulsion (which does not affect simulations of the two-quartet 15-TBA with just one internal cation) contributes to the unsatisfactory ΔG_fold_ of the three-quartet quadruplex. The magnitude of this effect is presently not clear. We note that the overestimation of inter-cation repulsion may be in future resolved by polarizable force fields (83, 84).

Further, it is possible that the force field understabilizes the propeller loops, as has been tentatively suggested in the past. Previous unbiased and temperature-accelerated MD studies reported notably fast unfolding (short lifetimes) for all-*anti* hairpins, triplexes, and G4s (the latter simulated in the absence of cations) with propeller loops. Longer lifetimes were observed for structures with propeller loops with at least one guanosine in *syn*, and structures with lateral and diagonal loops (26, 32, 34, 37, 85). This is consistent with the extraordinary instability of the parallel all-*anti* hairpin with the propeller loop seen in the present study which was for the first time quantified via calculated ΔG_fold_ (Table 3). However, it is unclear which parts of the force field would be responsible for the imbalanced description of the propeller loops. It does not appear to be caused by dihedral parametrizations, as differences in dihedral parameters of recent refinements of the AMBER force field are too subtle. For example, the ɑ/γ dihedral correction introduced in the OL21 DNA force-field update of the OL15 version improved B-DNA and Z-DNA descriptions (64, 86) but did not improve the stability of the G-hairpins simulated here (Supplementary data, Table S3).

### Exploration of the FES is very limited for such complex systems

Despite the predicted positive free energy of all target species, we can still characterize the folding events thanks to the metadynamics part of the simulation protocol, as the metadynamics flattens the FES along the used CV. The used ST-metaD approach combines well-tempered metadynamics with εRMSD CV biasing the simulations towards the target structures with REST2 replica-exchange protocol to accelerate sampling of the remaining degrees of freedom. Due to the richness of the studied systems’ FES, fully converged conformational ensembles are still out of our reach even for G-hairpins simulated with enhanced-sapling protocols. Isolated G-hairpins are unstable species per se, which further complicates their studies with unbiased MD simulations. The chosen CVs (i.e., the direction in which the bias is applied to guide the conformational sampling) may also affect the sampled states and folding pathways (87). The ST-metaD protocol applied here could capture the formation of very unstable species including folding a full three-quartet G4 and predicting their ΔG_fold_. It is however evident that the present simulations remain far from a quantitative convergence and the exploration of the unfolded conformational space is affected (restricted) by the used CV (see Supplementary data, Figure S14). We still suggest that the ST-metaD method is robust enough to qualitatively study molecules with complex FES and such elusive processes as G4 folding, though the latter is probably only possible in the vicinity of one specific fold (folding funnel) to which the CV is linked. We note that a similar technique of parallel tempering combined with metadynamics using 2D bias defined by εRMSD and RMSD was successfully used in the above-mentioned simulations of 15-TBA G4 (30, 35).

We suggest that the employed CV will affect the folding pathways of the full G4 more than those of isolated hairpins. At the same time, G4 folding landscapes are very complex and cannot be fully captured by a single simple CV. Despite that we have used the inter-tract εRMSD CV, biasing the εRMSD of all four G-tracts at the same time can excessively push them together. Thus, we made a partly biased simulation where only the first two G-tracts were biased by the CV. Comparison of the compactness of biased and unbiased segments of the (GGGA)_3_GGG chain reveals that the coil ensemble is readily sampled also in the unbiased segment, but its properties differ from the one under the bias (***Figure 4***a). Thus, while both biased and partly biased simulations support the existence of the coil ensemble, its sampling can be CV-dependent. This does not invalidate our six observed folding events (**Figure 3**a) but rather points to the possible existence of other alternative pathways that are poorly captured by the employed CV. Optimization of folding CVs is one of our future goals.

To follow on the choice of CV, Rg is commonly monitored in G4 simulations and sometimes even in experiments, as it is supposed to measure how well a given molecule is folded. The presumption is that high Rg corresponds to the unfolded states and the lowest Rg to the compact fully-folded G4 structure. However, the distribution of Rg observed in our folding simulations shows that the coiled ensemble has a lower Rg than the native G4 (***Figure 4*** and Supplementary data, Figure S8), as we have also suggested before with simulations of misfolded structures (88). Thus, Rg is not a sufficiently discriminative metric for folded versus unfolded states. In simulations, this issue could be alleviated by considering Rg of just the G4 core (without the loops), possibly combined with another CV but in experimental Rg measurements one cannot exclude parts of the molecule at wish.

## CONCLUSIONS

Characterization of G4 folding pathways including the transitory ensembles linking the main free energy states is an important puzzle piece helping to understand G4 properties and roles. As the characterization of short-living intermediates and transitory ensembles cannot be achieved by experimentation, it is a goal for complementary modeling and simulation studies (19, 20). Here, we report a series of all-atom enhanced-sampling folding simulations targeting the parallel three-quartet DNA G4 and several G-hairpin topologies.

The ST-metaD simulations suggest that the folding of parallel G4 proceeds by multi-pathway structuring of the quartets within a collapsed compact coil-like ensemble (perhaps resembling the molten globule state in protein folding, ***Figure 8***). The coil ensemble samples diverse guanine-guanine interactions and progresses toward the native G4 fold via multiple small incremental steps. The G4 structure thus emerges from the compact coil-like ensemble via numerous rearrangements. This is a quite different process compared to common literature folding models that proceed via very few simple intermediates, such as the Hoogsteen triplex. We hypothesize that the coil-like ensemble is important not only in the folding from unfolded states but also for transitions between different G4 topologies, if present on the landscape. With potential consequences for experimental measurements, we also show that the coil-like ensemble has a smaller radius of gyration than the native G4 (***Figure 4***) and therefore Rg might sometimes be ambiguous for following the G4 folding process.

Our study also brings significant methodological results. The ST-metaD simulation predicts a high positive ΔG_fold_ of +17 kcal/mol for the fully folded G4 (Table 2). To our best knowledge, it is the largest free energy discrepancy reported for nucleic acids simulations so far. Enhanced sampling simulations of the (GGGA)_3_GGG G4 structure thus can serve as a vital benchmark for DNA force field development. The free energy discrepancy is probably a result of several force field imbalances, as discussed above. Fortunately, folded G4 molecules can be quite reliably studied by standard simulations since they represent deep free energy basins on the FES and are separated from the unfolded ensemble by a large free energy barrier. Nevertheless, the present study gives a warning that the global description of the G4 folding landscapes is quite severely imbalanced. More studies will be needed to understand the origin of the imbalances and to see how (and if) they can be corrected.

Despite using a sophisticated enhanced sampling protocol, the simulations remain far from a rigorous convergence which would require obtaining equivalent sampling in all demuxed (continuous) trajectories. This has not been achieved even for the hairpin simulations. Still, we suggest that the G4 folding events reported above provide valid insights into the mechanism of G4 folding.

In summary, our work provides new atomistic insights into parallel G4 folding landscapes and the role of coil-like compacted ensembles. It also suggests that ST-metaD and similar simulation protocols could be used in further studies of G4 folding mechanisms and metastable ensembles, though realistically a fully converged sampling of the G4 landscapes is not within the reach of contemporary methods and computers. Finally, the current AMBER force fields severely underestimate the stability of folded parallel-stranded G4 with respect to the unfolded ensemble.

## DATA AVAILABILITY

The input files, stripped trajectory files (reactive trajectories and reference replicas for G4, and reactive trajectories, and reference replicas for selected hairpin simulations), bias files, and the ΔG_fold_ calculation protocol are available on Zenodo (DOI: 10.5281/zenodo.8247280) and on GitHub (www.github.com/sponerlab/G4_folding_parallel). PLUMED input files are available in the plumed-nest.org repository ([plumID:23.033](https://www.plumed-nest.org/eggs/23/033/)).

## ACKNOWLEDGEMENTS

We acknowledge the computational resources provided by the Ministry of Education, Youth and Sports of the Czech Republic through the e-INFRA CZ [ID:90254].

