## Supplementary material for "Parallel G-quadruplex folds via multiple paths involving G-tract stacking and structuring from coil ensemble"

Other supplementary files (structures, simulation setting files, $\Delta G_{fold}$ calculation protocol) are provided in a separate .zip archive and are also available at GitHub repository www.github.com/sponerlab/G4_folding_parallel. Trajectory data and bias files (HILLS) are available in the Zenodo database (DOI: 10.5281/zenodo.8247280), PLUMED input files are also available at plumed-nest.org, plumID:23.033.

**Table S1**: Folding free energies and numbers of reactive trajectories in individual simulations

| **n.** | ref. structure | target loop topology, *syn-anti* G-tract pattern, groove width[a] | force field | no. of replicas | length [$\mu$s] | additional settings[b] | $\Delta G_{fold}$ of the target biased state [kcal/mol] | no. of replicas with folding events[c] (proportionally to no. of replicas) |
|---|---|---|---|---|---|---|---|---|
| | | | | **GGGAGGGAGGGAGGG** | | | | |
| **1** | G4 | propeller, all-*anti*, m | OL21 | 16 | 3.7 | inter-tract CV | 16.9 | 4 (25%) |
| **2** | | | OL21 | 16 | 0.5 | inter-tract CV | - | 2 (12%) |
| **3** | hairpin | propeller, all-*anti*, m | OL21 | 16 | 2 | inter-tract CV, only first two G-tracts biased | 9.1 | 11 (69%) |
| **4** | G4 | propeller, all-*anti*, m | OL21 | 16 | 2 | inter-tract CV, abasic loops | 9.3 | 7 (44%) |
| **5** | G4 | propeller, all-*anti*, m | OL21 | 16 | 2 | inter-tract CV, gHBfix (2 kcal/mol) | 18.1 | 2 (12%) |
| | | | | **GGGAGGGAGGG** | | | | |
| **6** | triplex | propeller, all-*anti*, m | OL21 | 16 | 3 | | 20.1 | 12 (75%) |
| | | | | **GGGAGGG** | | | | |
| **7** | hairpin | propeller, *aaa-aaa*, m | OL15 | 12 | 4.5 | | 18.7 | 10 (84%) |
| **8** | | | OL21 | 12 | 5 | | 16.9 | 9 (75%) |
| | | | | **GGGTTAGGG** | | | | |
| **9** | hairpin | propeller, *aaa-aaa*, m | OL15 | 16 | 5 | | 11.7 | 14 (88%) |
| **10** | | | OL15 | 16 | 5 | | 12.5 | 15 (94%) |

| | | | | | | | |
|---|---|---|---|---|---|---|---|
| **11** | | OL15 | 12 | 5 | | 11.2 | 12 (100%) |
| **12** | | OL15 | 16 | 5.5 | inter-tract CV | 12.4 | 14 (88%) |
| **13** | | OL15 | 12 | 5 | inter-tract CV, structure-specific HBfix (6x2 kcal/mol) | 3.3 | 12 (100%) |
| **14** | | OL15 | 12 | 5 | inter-tract CV, structure-specific HBfix (6x2 kcal/mol) | 2.7 | 9 (75%) |
| **15** | | OL15 | 12 | 5 | inter-tract CV, gHBfix (2 kcal/mol) | 13.0 | 7 (58%) |
| **16** | | OL15 | 12 | 5 | inter-tract CV, gHBfix (2 kcal/mol) | 14.5 | |
| **17** | | OL15 | 12 | 4 | abasic loop | 10.1 | 12 (100%) |
| **18** | propeller, *saa-saa*, m | OL15 | 16 | 5 | | 8.4 | 16 (100%) |
| **19** | propeller, *saa-saa*, m | OL15 | | 4 | | 7.9 | 15 (94%) |
| **20** | lateral, *saa-ssa*, W | OL15 | 16 | 4.5 | | 6.4 | 14 (88%) |
| **21** | | OL15 | 12 | 5 | | 5.3 | 10 (86%) |
| **22** | | OL15 | 16 | 5 | 266-700 K temperature range | 5.6 | 16 (100%) |
| **23** | lateral, *ssa-saa*, W | OL15 | 12 | 4.8 | | 5.1 | 10 (84%) |
| **24** | lateral, *saa-ssa*, n | OL15 | 12 | 4.8 | | 4.2 | 10 (84%) |

| | | | | | | | |
|---|---|---|---|---|---|---|---|
| **25** | lateral, *ssa-saa*, n | OL15 | 12 | 5 | | 5.7 | 16 (100%) |

[a] for structures, see the main text, Figure 2.

[b] unless stated otherwise, simple εRMSD was used as a CV, effective temperature range of 298-500 K was applied, and no (g)HBfix was used.

[c] regardless the number of folding events.

**Table S2**: Populations of stacked G-tracts in the previously published REST2 simulations run with torsional restraints applied on Gs to remain in *anti,* no metadynamics bias was applied.

| | stacked G-track population [%][a] | | |
|---|---|---|---|
| system | G-tract 1 | G-tract 2 | G-tract 3 |
| GGGAGGG | 76 | 73 | - |
| GGGTTAGGG | 40 | 40 | - |
| GGGAGGGAGGG | 34 | 68 | 56 |

[a] cutoff for considering G-tract stacked is 80 Å$^2$ interaction surface of the three bases.

**Table S3**: OL15 vs. OL21 folding free energies

| simulation, force field | $\Delta G_{fold}$ [kcal/mol] | $\Delta G_{fold}$ reweight to OL21 [kcal/mol] | $\Delta G_{fold}$ reweight to OL15 [kcal/mol] |
|---|---|---|---|
| GGGTTAGGG, *aaa-aaa*, m, inter-tract CV, OL21 | 12.4 | 12.4 | - |
| GGGAGGG, OL15 | 18.8 | 18.6 | - |
| GGGAGGG, OL21 | 17.0 | - | 16.8 |

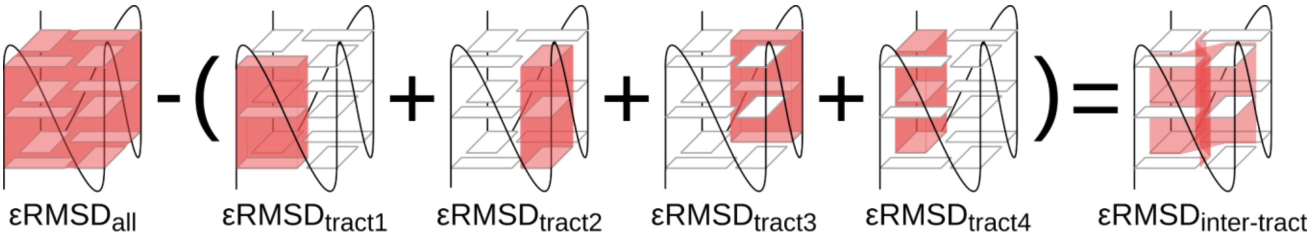


**Figure S1**: Schematic representation of the inter-tract εRMSD CV for the full G4.

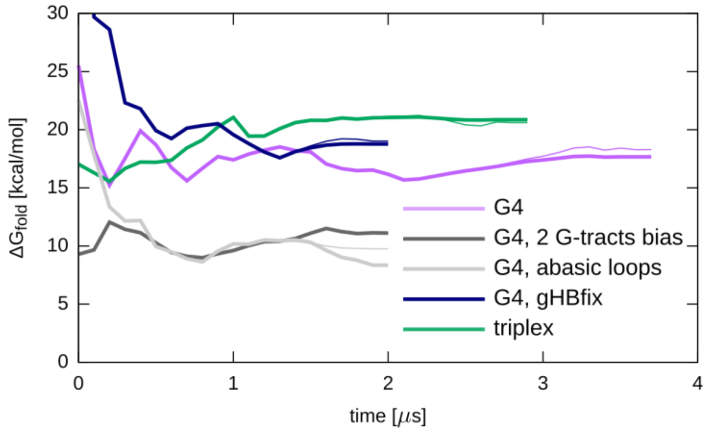


**Figure S2**: Bias convergence in G4 and triplex simulations calculated from the reference (unbiased) replica. Thick lines show convergence of averaged bias.

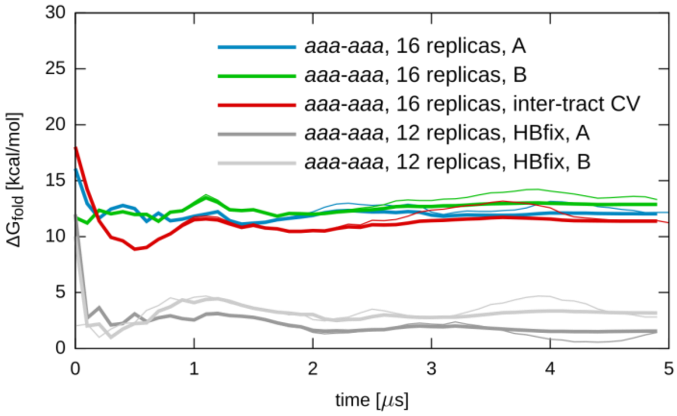


**Figure S3**: Bias convergence in selected GGGTTAGGG simulations. Thick lines show convergence of averaged bias.

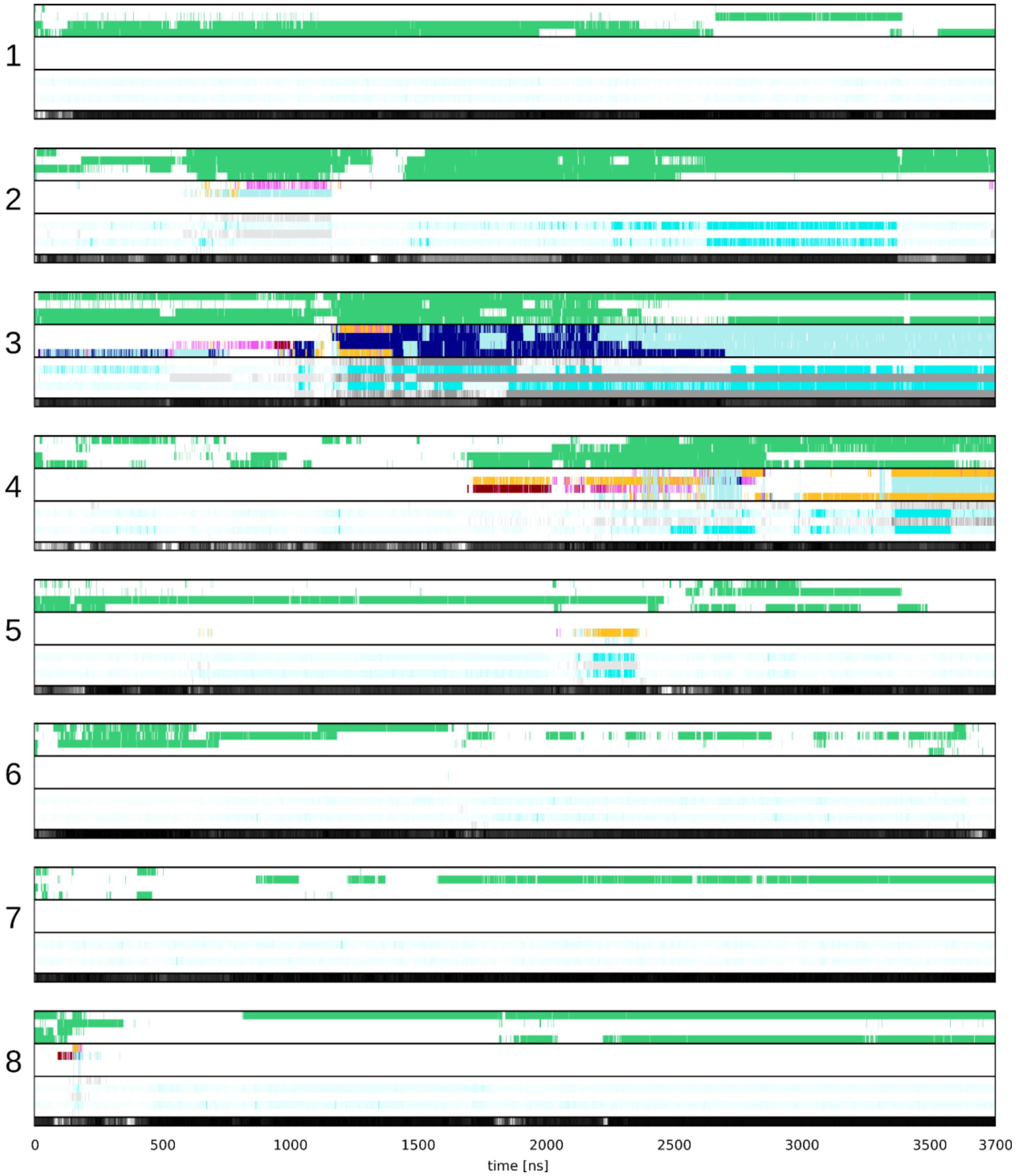


**Figure S4**: Demuxed trajectories 1-8 of G4 simulation (simulation no 1., see Table S1). For legend, see the main text Figure 3.

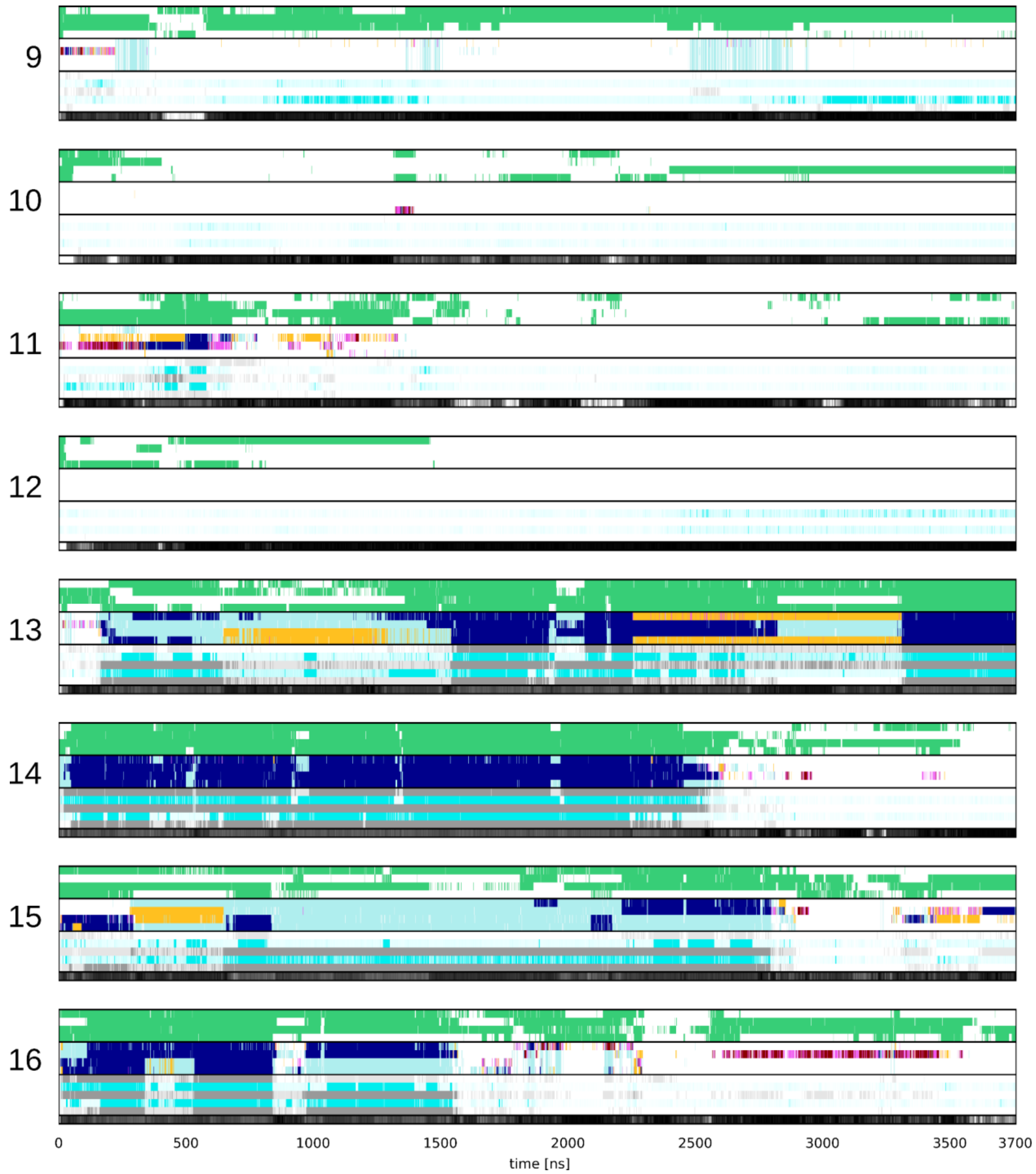


**Figure S5**: Demuxed trajectories 9-16 of G4 simulation (simulation no 1., see Table S1). For legend, see the main text Figure 3.

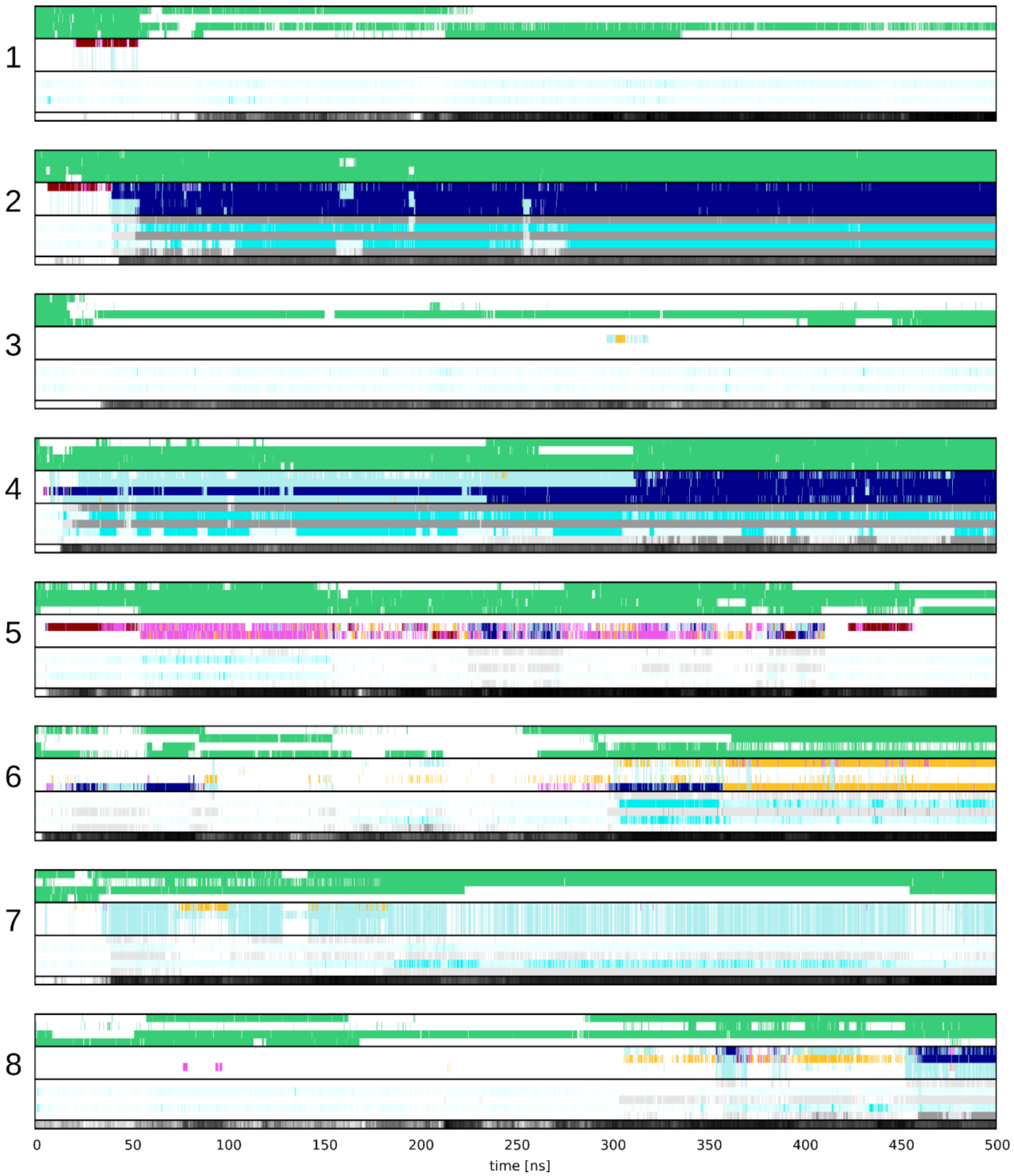


**Figure S6**: Demuxed trajectories 1-8 of G4 simulation (simulation no. 2, see Table S1). For legend, see the main text Figure 3.

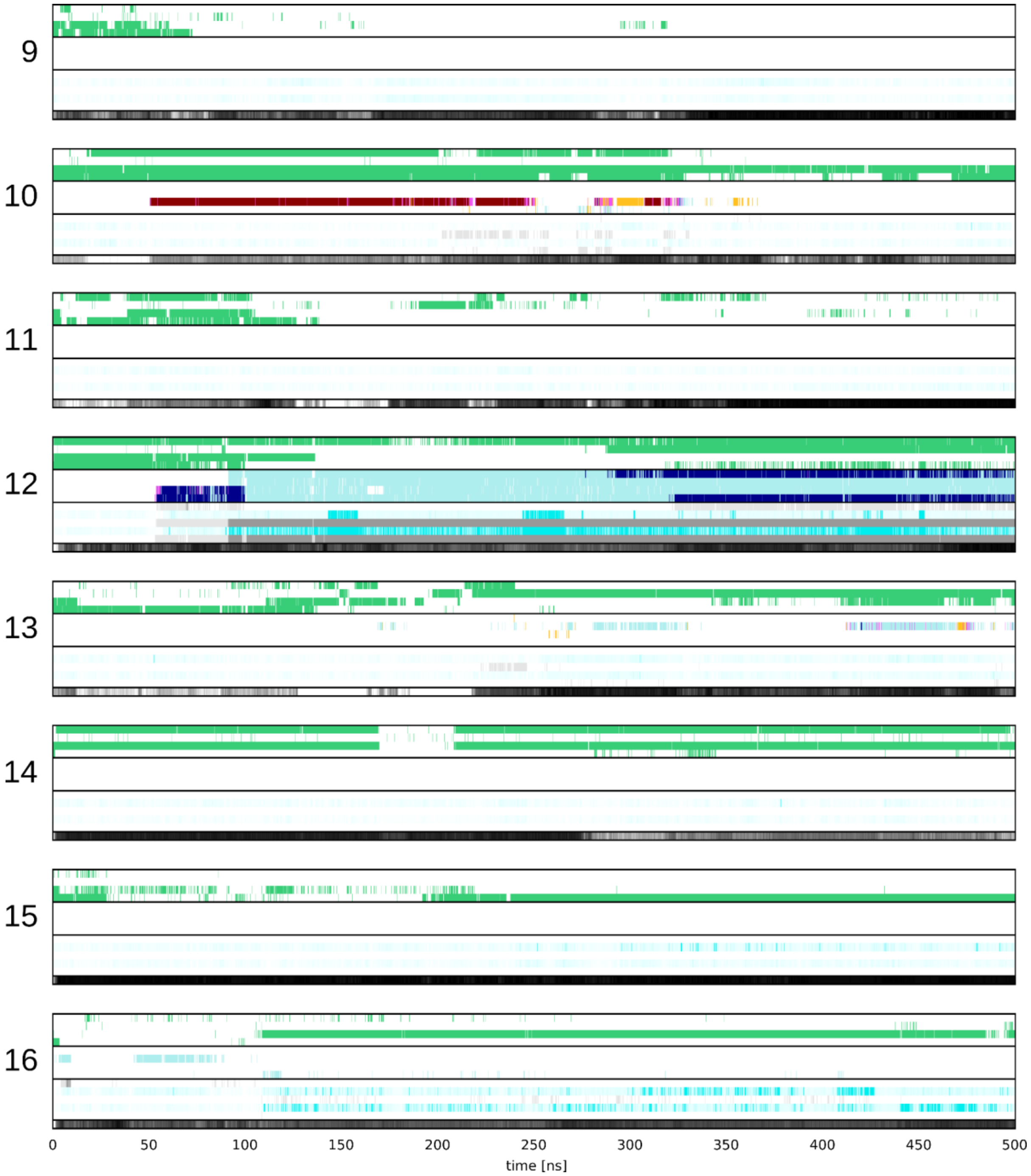


**Figure S7**: Demuxed trajectories 9-16 of G4 simulation (simulation no. 2, see Table S1). For legend, see the main text Figure 3.

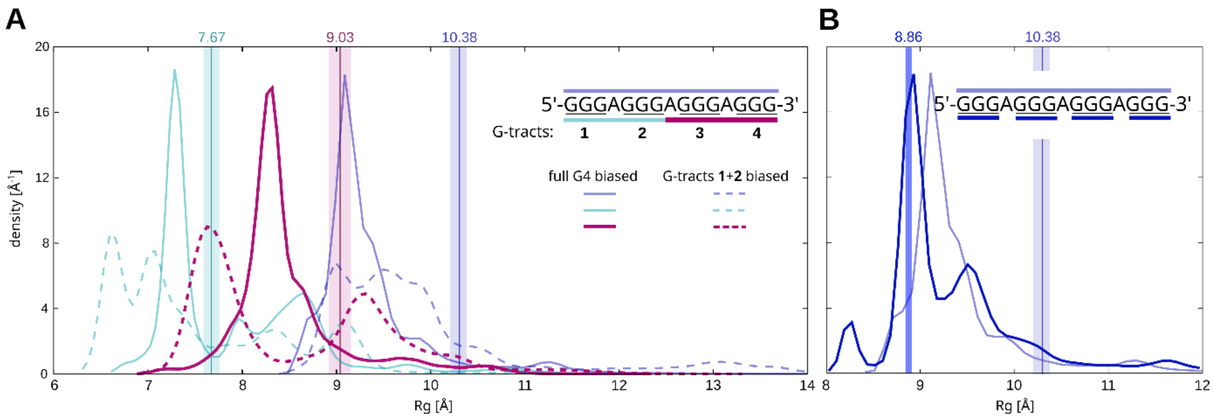


**Figure S8**: Radii of gyration in simulations of $(GGGA)_3GGG$ sequence, reweighted densities. See the main text Figure 4 for details.

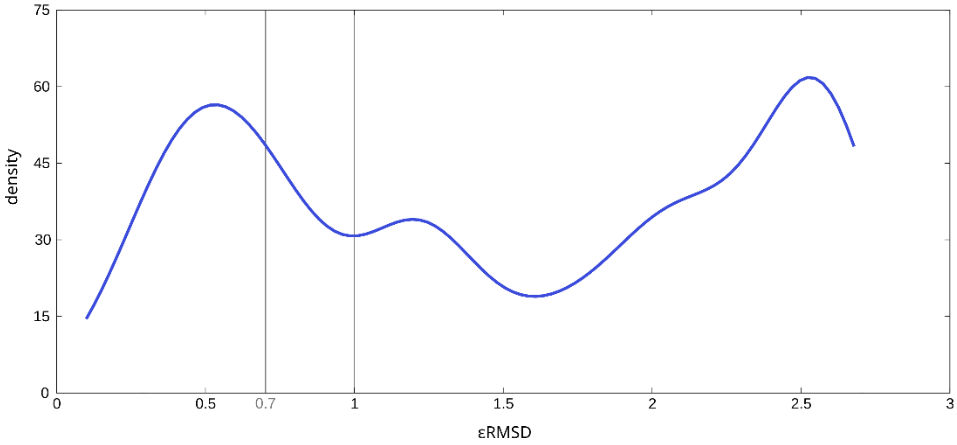


**Figure S9**: εRMSD distribution for structures with Rg (calculated only for Gs) between 8.8 and 8.9 Å. εRMSD is calculated with respect to the folded G4. Data are from the reference replica of simulation no. 1.

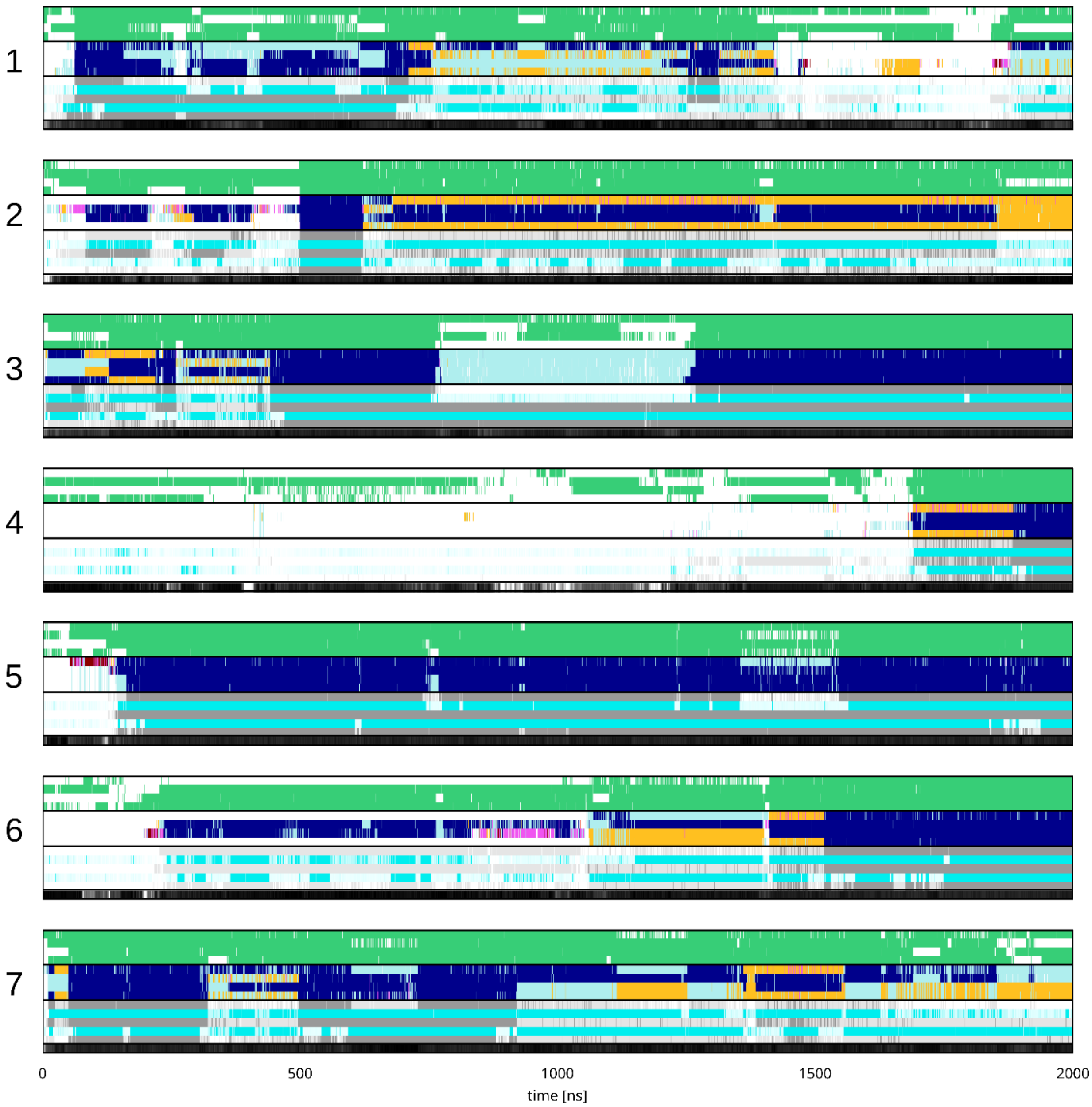


**Figure S10**: Demuxed trajectories with folding events in G4 simulation with abasic loops (simulation no. 4, see Table S1). For legend, see the main text Figure 3.

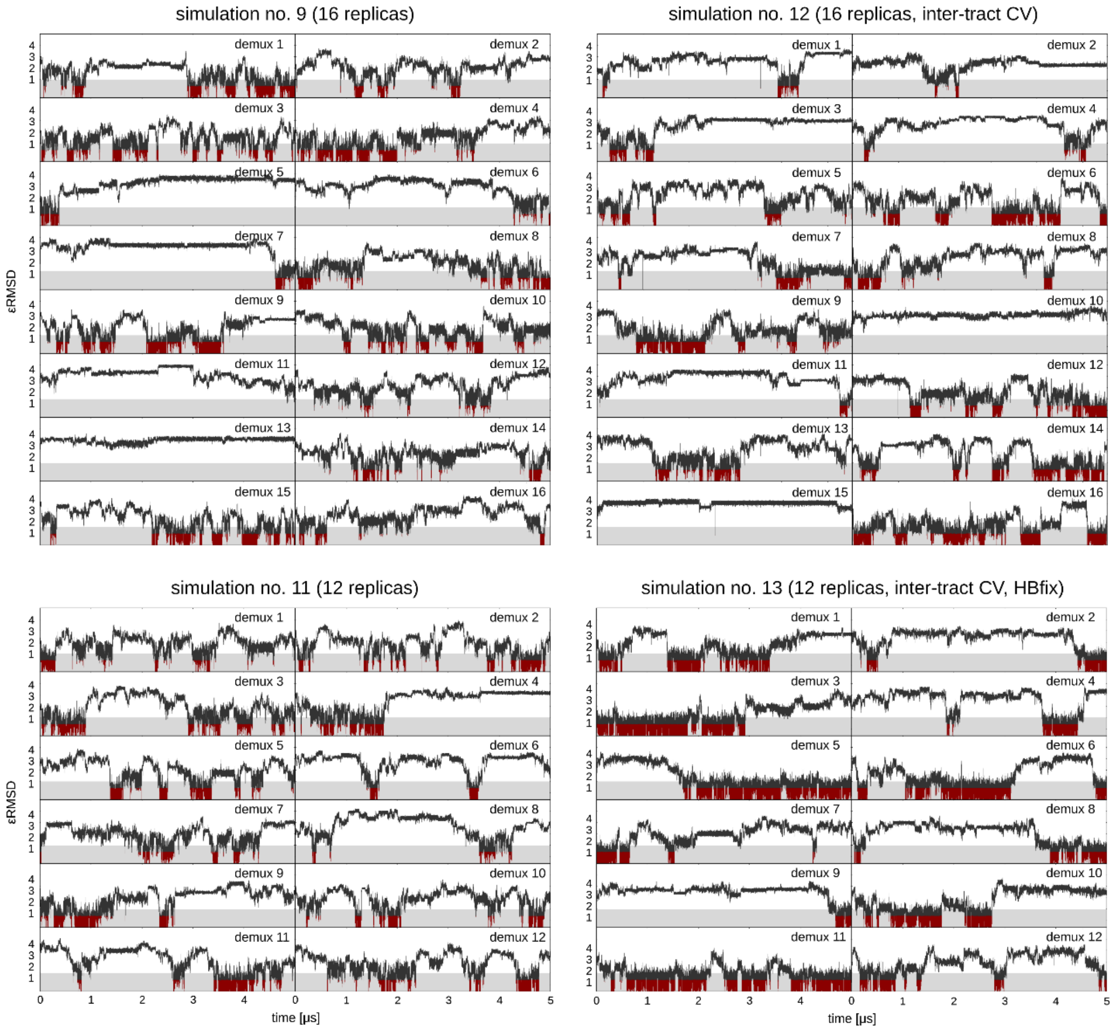


**Figure S11**: Sampling of GGGTTAGGG folding events in four selected demuxed trajectories with different simulation settings. Gray areas mark εRMSD values between 0 and 1 and trajectory parts with εRMSD < 0.7 are in red.

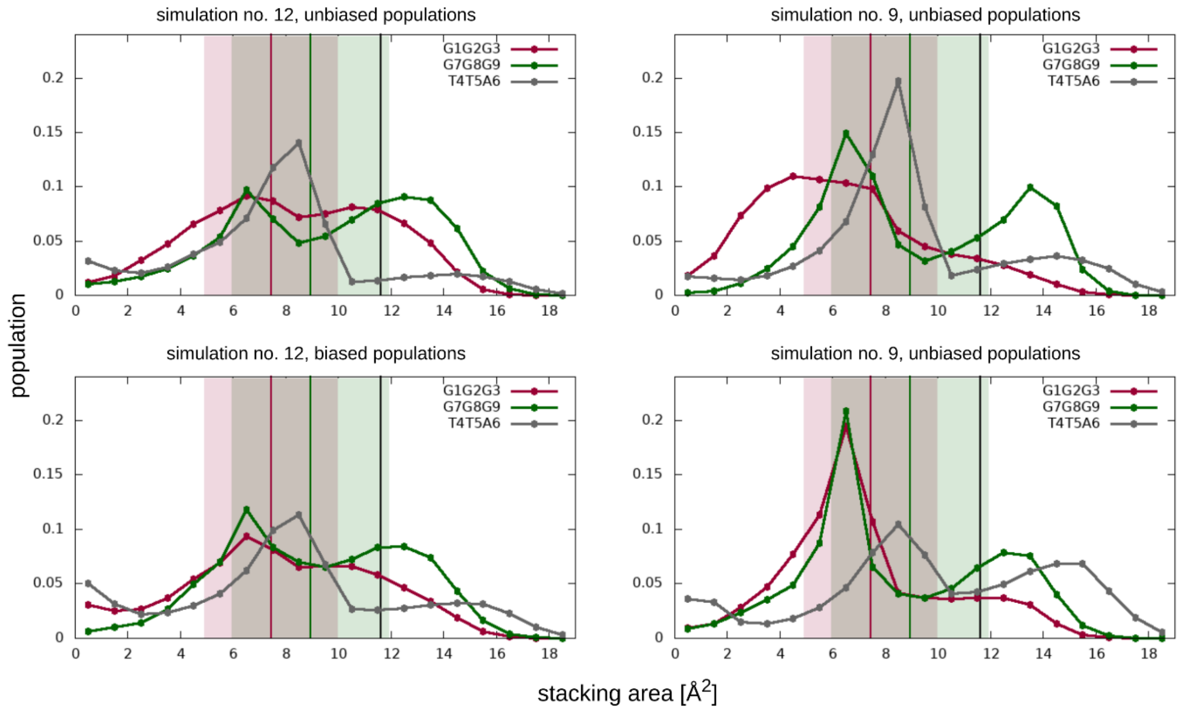


**Figure S12**: Stacking surface area in reference replicas of GGGTTAGGG *aaa-aaa* simulations with the inter-tract εRMSD CV (left, simulation no. 12), and full εRMSD CV (right, simulation no. 9). The colored vertical lines show average values from the folded ensemble (εRMSD < 0.7) and the solid background the respective standard deviations. The black vertical line corresponds to the GpGpG stack in B-DNA. Stacking area was calculated with DSSR-x3DNA (1).

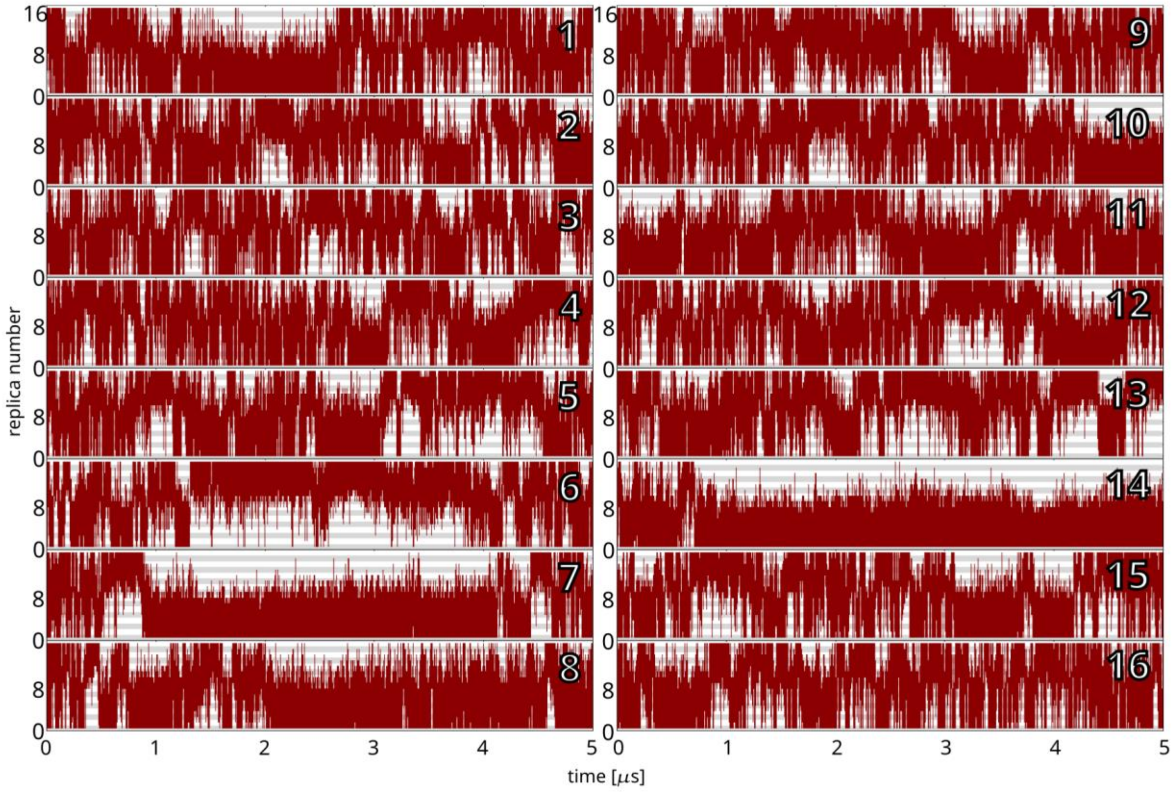


**Figure S13**: Replica ladder travel by 16 demuxed trajectories in one GGGTTAGGG simulation targeting the propeller/*aaa-aaa*/m hairpin topology.

### Ensemble overlap for simulations targeting different hairpin topologies

To visualize the overlap between the ensembles sampled in different simulations, we compare reference replica ensembles of simulations biasing the *aaa-aaa* parallel and *saa-ssa*, w, antiparallel hairpins (Figure S14). We chose these two target topologies for the analysis since they are very different. Simulation settings were identical, except for the target CV structure. Comparison of simulation ensembles is based on εRMSD clustering performed with an in-house modified version of Barnaba (2). Reference replica trajectories from both simulations were merged, and the clustering was performed on the merged ensemble. The εRMSD cutoff was set to 0.7 and the sensitivity to 0.001. With these settings, 82% of the frames were assigned to 125 clusters. 32 clusters were overlapping between the two simulations with εRMSDs from the reference structures around 1.5 to 2.2. In total, 5% of frames per trajectory overlapped. The actual overlap could be higher due to potential overlaps among the unassigned frames and the fact that some of the 125 different clusters could be similar. The median value of the lowest RMSD between two different clusters was 6.8 Å, and the minimum was 5.3 Å, considering the full GGGTTAGGG sequence for clustering. When the loop was omitted, the median value decreased to 3.1 Å, the minimum to 1.3 Å, and there were 21 clusters (out of 125) with RMSD < 2.0 Å to some other cluster. For the 40 most-populated ones (as shown in Figure S12), the clusters with RMSD < 2.0 Å were always significantly sampled only in the same trajectory.

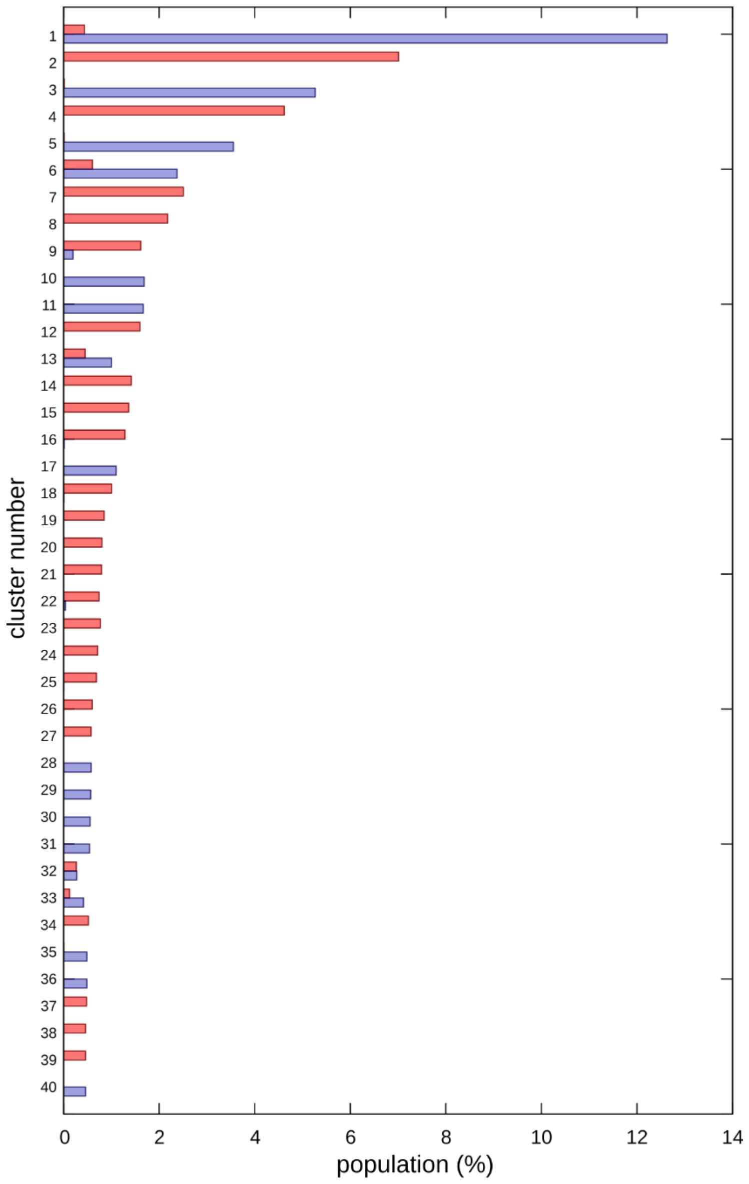


**Figure S14**: Cluster analysis on the two ensembles from simulations biased towards different hairpin topologies. Clusters are numbered from the most populated (1) to the least. Only the first 40 clusters are shown.
